# MOSAIC: Intra-tumoral heterogeneity characterization through large-scale spatial and cell-resolved multi-omics profiling

**DOI:** 10.1101/2025.05.15.654189

**Authors:** MOSAIC Consortium, Caroline Hoffmann

## Abstract

Precision oncology remains challenging due to gaps in understanding tumor biology and immunology, as well as the scarcity of consistent data across large cohorts. Here, we introduce MOSAIC (Multi-Omics Spatial Atlas in Cancer), a multi-center clinical Omics study designed to systematically profile thousands of cancer samples, with a focus on spatial and single-cell data across multiple tumor types. MOSAIC exploits recent technological advances to integrate spatial and single-cell data with complementary data modalities, including hematoxylin-eosin histology scans, bulk RNA and exome sequences, to generate comprehensive representations of cancer histology, genomics, and transcriptomics. MOSAIC also collects extensive and curated clinical information to ensure that patients meet the precise inclusion criteria for each cohort. The consortium aims to integrate all data modalities using artificial intelligence and other computational approaches in order to identify clinically-relevant biomarkers and cancer subtypes. This paper outlines the core objectives of MOSAIC, its potential impact, early proofs-of-concept, design, and experimental considerations. Additionally, we introduce the MOSAIC Window initiative, featuring the first released dataset from 60 patients, offering a glimpse into the project’s groundbreaking potential. Using four selected patients, we demonstrate the power of multi-omic approaches to analyze and interpret intra-tumoral variability within and across patients, providing insights on specific drug sensitivity of cancer subpopulations and revealing potential impacts on therapeutic recommendations.

## Introduction

Cancer research has made tremendous advances over the past 50 years, making many cases treatable with surgery, radiation, chemotherapies, and targeted drugs aimed at cancer driver genes. The past ten years particularly have seen the introduction of immune checkpoint inhibitors (ICI) that can unleash the host’s formidable arsenal of oncolytic lymphocytes in cases where cancers develop immune resistance (Shiravand et al 2022). However, despite this progress, too many mid- to late-stage cancer patients remain unresponsive to therapy. For example, PD1/PDL-1 inhibitors, which are currently the best available treatment for many cancers, have <50% response rates less at best, and are less than 20% effective or have failed to demonstrate any improvement in many cancer indications (Chen et al 2021, Yamaguchi etal 2024). This high rate of refractory tumors arises from a variety of causes, but a major component is limited understanding of complex tumor-immune interactions. In essence, patients exhibit a wide variety of immune activity in the presence of tumors, which if properly described, would enable us to discover new drug targets, and redirect nonresponsive patients away from therapies that simply won’t work for them. For this to be achieved, it is essential to consider each cancer indication as comprising multiple subtypes, with distinct tumor-immune biological mechanisms (Wahida et al 2023).

Precision oncology consists of moving from clinical diagnoses to biologically distinct subtypes, which may or not be common across different cancer types, with the end goal of personalized therapies for each group of patients (Garraway et al, J Clin Onc, 2013). This strategy is enabled by recent advances in technologies for medical imaging, molecular profiling, and artificial intelligence (AI). Spatially resolved molecular profiling (“spatial omicsˮ) is a recent technology that harnesses all three of these advances (Ståhl et al, Science, 2016; Rao et al, Nature, 2021; Zeng et al. 2022). As cancer is driven by clonal diversity as well as localized interactions between tumor cells, immune and non-immune stromal components, spatial omics have emerged as a key methodology to reveal specific cell types and pathways that control those tumor microenvironment (TME) crosstalks (Hunter et al, Nat Comm, 2021; Alon et al, Science, 2021). Spatial omics are poised to transform our understanding of cancer and its interaction with the immune system, involving both innate and adaptive immunity (Moses er Pachter., 2022; Marx, 2021; Binnewies et al, Nat Med, 2018). Due to their high dimensionality, sparsity, and challenging signal-to-noise ratios, current spatial transcriptome technologies greatly benefit from integration with complementary data modalities at single-cell resolution, enhancing the quality and depth of information (Schmauch et al, bioRxiv, 2024). Incorporating spatial omics with single-cell sequencing can therefore create synergy by mapping the spatial relationships and potential interactions among various cell types within the TME (Andersson et al., Communications Biology 2020). Therefore, the multi-omics approach can also offer additional insights for therapeutic targeting (Lyubetskaya et al., 2022). Advances in AI applied to various aspects of ST analysis, including identification of spatially variable genes, cell-cell communication and cell type deconvolution, hold significant promise for driving biological discoveries, such as the identification of new cancer subtypes, predictive biomarkers of treatment response/resistance (Jin et al., Mol Cancer 2024; Ren et al., Nat Commun 2023), or as-yet unknown spatial structure of tumor-immune interactions (Liu et al., J Hepatol, 2023). However, advancements in AI methodologies for developing such approaches have so far been hindered by the small sample sizes typical of spatial omics studies. Additionally, variations in protocols and technologies make the integration of disparate spatial omics datasets challenging.

Here we present MOSAIC (Multi-Omics Spatial Atlas in Cancer) - a collaborative initiative across industry and top cancer centers to build the largest collection of spatial omics data in oncology. By integrating comprehensive clinical annotations with advanced deep profiling techniques, the project aims to uncover novel cancer subtypes and identify key drug targets and biomarkers. MOSAIC is generating multimodal data targeting patients in at least nine tumor types: Non Small Cell Lung Cancer (NSCLC), Ovarian Cancer (OV), Bladder Urothelial Carcinoma (BLCA), Mesothelioma (MESO), Glioblastoma (GBM), Breast Invasive Carcinoma (BC), Diffuse Large B Cell Lymphoma (DLBCL), Head and Neck Squamous Cell Carcinoma (HNSCC), and Pancreatic adenocarcinoma (PAAD). As of April 2025, more than 1900 patients have been included in the study.

## MOSAIC Consortium

### MOSAIC Data Modalities

MOSAIC studies the biology of tumor samples from patients with various clinical conditions (Figure 1A; Methods A.1.), by incorporating six distinct data modalities (Figure 1B, Methods A.2.) from medical records and FFPE specimens: i) Comprehensive clinical data, curated to ensure that patients meet the specific inclusion criteria for each predefined cohort; ii) Hematoxylin and Eosin (H&E) stained microscopic images, obtained with standard site-specific staining protocols; iii) Spatial Transcriptomics (ST) generated using the Visium Cytassist standard protocol from 10X Genomics; iv) Single-Nuclei RNA transcriptomics (snRNAseq), utilizing probe-based technology from Chromium Flex (10x Genomics); v) Bulk RNA-sequencing (bRNAseq), providing whole transcriptome profiling; vi) Whole Exome Sequencing (WES), offering detailed genomic insights.

**Figure 1.**
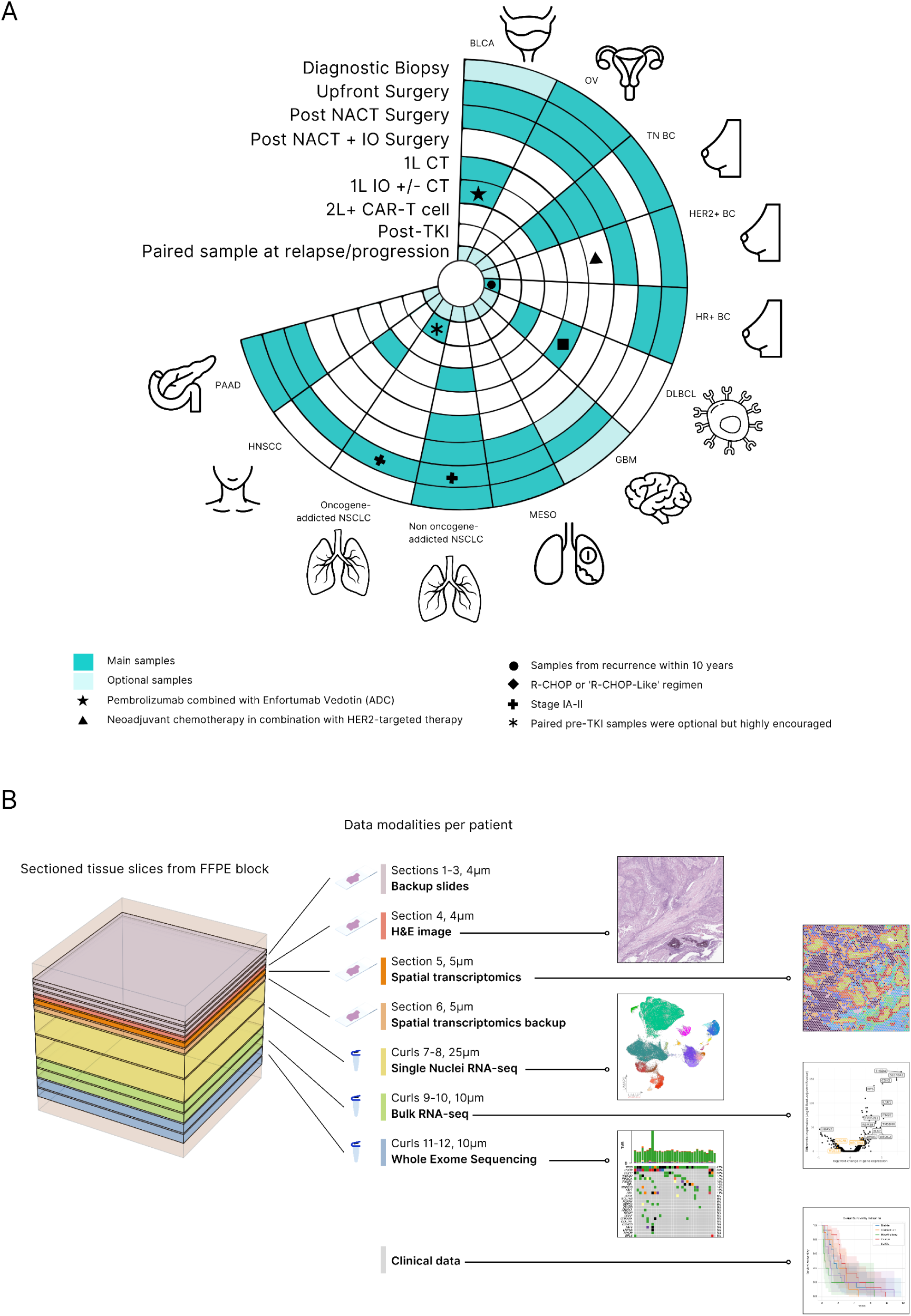
Overview of MOSAIC cohorts and data modalities. (A) Patient Cohorts and Clinical Settings. (B) Schematic Representation of Data Modalities. Each FFPE block is subjected to sectioning, where six consecutive sections are made for H&E and Visium assays, and as back-up sections. Following the preparation of sections, six curls are cut from the remaining FFPE block for snRNA, bRNAseq and WES analysis (Two curls per modality). Abbreviations: H&E, hematoxylin & eosin; snRNAseq, single-nuclei RNA sequencing; bulkRNAseq, bulk RNA sequencing; WES, whole-exome sequencing; NACT, neoadjuvant chemotherapy; IO, immuno-oncology; CT, chemotherapy; CAR T-cell, chimeric antigen receptor T-cell therapy; TKI, tyrosine-kinase Inhibitor; R-CHOP, rituximab/cyclophosphamide/hydroxydaunorubicin/oncovin/prednisone; BLCA, Bladder Urothelial Carcinoma; OV, Ovarian serous adenocarcinoma; TN BC, triple-negative breast cancer; HER2+ BC, HER2-positive breast cancer; HR+ BC, hormone-receptor positive breast cancer; DLBCL, diffuse large B-cell lymphoma; GBM, glioblastoma multiforme; MESO, mesothelioma; NSCLC, non small-cell lung carcinoma; HNSCC, head & neck squamous-cell carcinoma; PAAD, Pancreatic adenocarcinoma.

One of the key advantages of the MOSAIC study is the centralization of all the data collected and generated in Owkin central data platform, *Owkin K*. Each data modality is curated and processed separately through a dedicated processing workflow relying on state of the art methods (Figure 2; Supplementary Figure 1; Methods A.3.). In addition, clinical data are curated and mapped to a common vocabulary to ensure a consistency between centers and tumor type.

**Figure 2.**
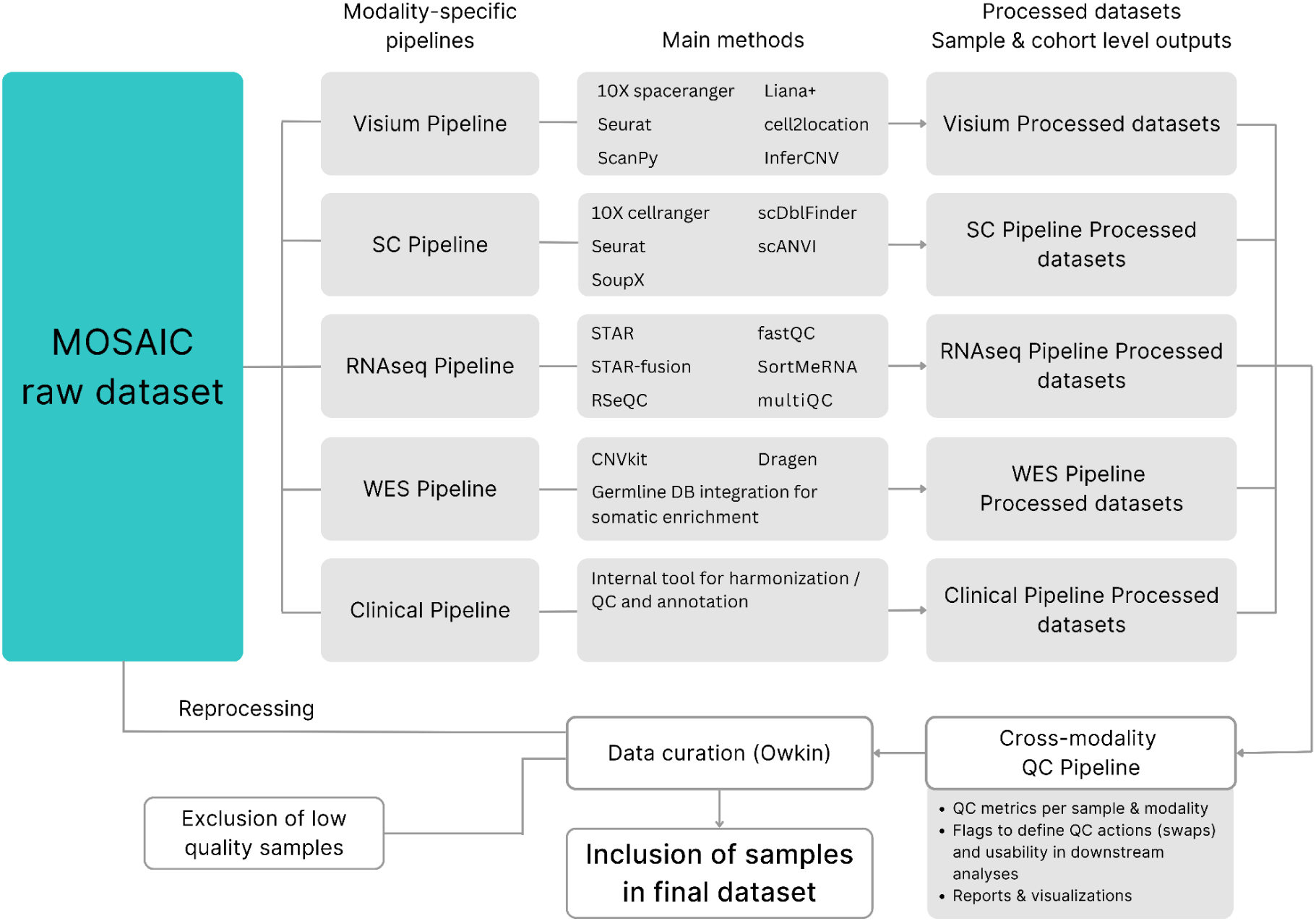
Overview of processing and data quality steps for all MOSAIC modalities. This schematic summarizes the end-to-end processing of raw data from MOSAIC patients across five modalities. For each sample block, data resulting from one modality is processed separately through a dedicated processing workflow relying on state of the art methods. These pipelines process each dataset (one dataset per center and cancer type) to produce gene expression quantification (single-nuclei RNAseq, bulk RNAseq, spatial-RNAseq), fusion calling (bulk RNAseq), mutation and copy-number calling (Whole-Exome Sequencing) and H&E tile level features (both for Whole Slide Images and spot-aligned spatial transcriptomics tiles). In addition, clinical data are curated to ensure a consistency between centers and indications with respect to inclusion criteria and flag any deviations. Abbreviations: SC, single-cell (referring to single-nuclei sequencing); RNAseq, RNA sequencing; WES, whole-exome sequencing; QC, quality control

### Consortium setup: Founding partners, regulatory framework, ethics committees and governance

The MOSAIC consortium is comprised of Owkin France as the sponsor of the study and five Founding Partners (FPs): Lausanne University Hospital, known as Centre Hospitalier Universitaire Vaudois (CHUV) in Lausanne, Switzerland; Charité – Universitätsmedizin Berlin, located in Berlin, Germany; The University Hospital Erlangen (UKER), part of Friedrich-Alexander-Universität (FAU) Erlangen-Nürnberg, is located in Erlangen, Germany; Institut Gustave Roussy (GR), located in Villejuif near Paris, France; The University of Pittsburgh, located in Pittsburgh, USA. The FPs are responsible for a significant part of the data generation workflow. Data is generated in dedicated MOSAIC laboratories and teams at each FP site, including standardized equipment across all locations. All relevant Institutional Review Boards (IRB) and Independent Ethics Committees (IEC) approved the clinical protocol (Supplementary Figure 2). The MOSAIC data generation process follows Standard Operating Procedures (SOPs) for each workflow step and relies on collaboration with key technology partners and sequencing facilities: 10x Genomics, CeGaT, GENEWIZ by Azenta Life Sciences, and Viedoc. A Joint Steering Committee (JSC) composed of members from Owkin and all FPs governs the consortium, overseeing MOSAIC resources and ensuring the study’s execution complies with all applicable regulations.

### Cohort mapping

The MOSAIC consortium has initially approved analysis into nine tumor types, but retains the possibility to extend the study to other solid tumors. Cohort selection by lead clinicians and pathologists per FP was based on a combination of factors: i) unmet medical need; iii) associated scientific questions; and iii) availability of relevant samples. Cohorts were cataloged according to tumor type, in subsequent “Cohort Proposalˮ and “Cohort Mappingˮ phases (Figure 1A; Supplementary Figure 3A-C; Methods A.1). To ensure sufficient statistical power for downstream analyses, we aimed for a minimum of 80 samples per cohort, while maximizing the synergy and contributions of all FPs. However, smaller cohorts are allowed for rare diseases or novel treatments. The catalog of cohorts is available in Table B.

## MOSAIC Window: a data release initiative

Here we announce the launch of MOSAIC Window, an initiative to make a MOSAIC dataset publicly accessible. The dataset comprises 60 cancer cases: BLCA (15), OV (15), GBM (10), DLBCL (10), and MESO (10). BLCA cases derived from upfront cystectomies, with most receiving complete lymph node dissection and some treated with platinum/gemcitabine or ICIs at relapse. OV samples include baseline, post-NACT primary tumors, and relapse lesions, with treatments including Bevacizumab, PARP inhibitors, and ICIs. GBM and DLBCL samples were obtained at baseline with standard therapies (Temozolomide-based chemoradiotherapy for GBM and R-CHOP for DLBCL)-supplemented by targeted interventions at relapse. MESO cases were baseline biopsies and surgeries and one post-NACT specimen. See Supplementary Table C and Methods B.1. for more details.

MOSAIC Window includes the six MOSAIC data modalities. Data was processed, curated and reviewed with strict quality control metrics. For ST, samples had a median UMI (Unique Molecular Identifier) per spot of 13 136 UMI and an average of 94% of reads mapped confidently to probe set. For snRNAseq, we had an average of 8 128 cells per sample and an average of 17 030 genes detected per sample. Additional data quality metrics for all modalities are reported in Supplementary Table D. Here, we present biological insights from MOSAIC Window cases (Methods B.2. for details on the data analysis).

The dataset is hosted on the European Genome-Phenome Archive (EGA) and researchers from both academic and industrial sectors can apply for access by submitting a detailed research proposal. A Data Access Committee (DAC), consisting of representatives from Owkin and FPs, evaluates the requests and grants access to the dataset.

### Results: deep dive on four MOSAIC Window cases

Cancer deep phenotyping with the six MOSAIC data modalities including high quality and detailed clinical data allows for unprecedented analysis of clinical cases and the corresponding cancer biology, with a potential impact on cancer therapy. Here, we present 4 detailed cases of the MOSAIC Window cohort in which the cancer heterogeneity was deciphered by multimodal data integration. We focused the final interpretation of each case on the potential therapeutic impact of the findings.

#### Case 1 - (MW_M_051)

A 47-year-old male patient with a history of active alcohol consumption and smoking was diagnosed with cT4NxM0 diffuse epithelioid mesothelioma, with no evidence of metastatic disease. He received four cycles of neoadjuvant cisplatin and pemetrexed chemotherapy, achieving a partial response. Four months after diagnosis, he underwent pleurectomy/decortication with multiple lymph node resections. Surgical pathology revealed ypT4N1 disease with a pleomorphic epithelioid subtype exhibiting over 50% solid growth, a high mitotic rate, and nuclear grade 3. He received one cycle of adjuvant pemetrexed, but treatment was discontinued due to toxicity. Less than six months after diagnosis, imaging studies revealed metastatic relapse involving a distant lymph node and the liver, and the patient died from disease progression within one week of this finding.

This post-chemotherapy tumor exhibited two distinct histological regions with different cellular compositions. One region is characterized by tumor cells with abundant cytoplasm, large round nuclei, and prominent nucleoli; these features are characteristic of the epithelioid pattern. In contrast, the other area shows abundant inflammation, with tumoral cells presenting marked pleomorphism, with occasional cells displaying a polygonal morphology and a loss of cohesiveness compared to the previous area; these findings suggest a transition zone observed in the sarcomatoid differentiation of mesothelioma which may have been induced by the treatment. (Figure 3A).

**Figure 3.**
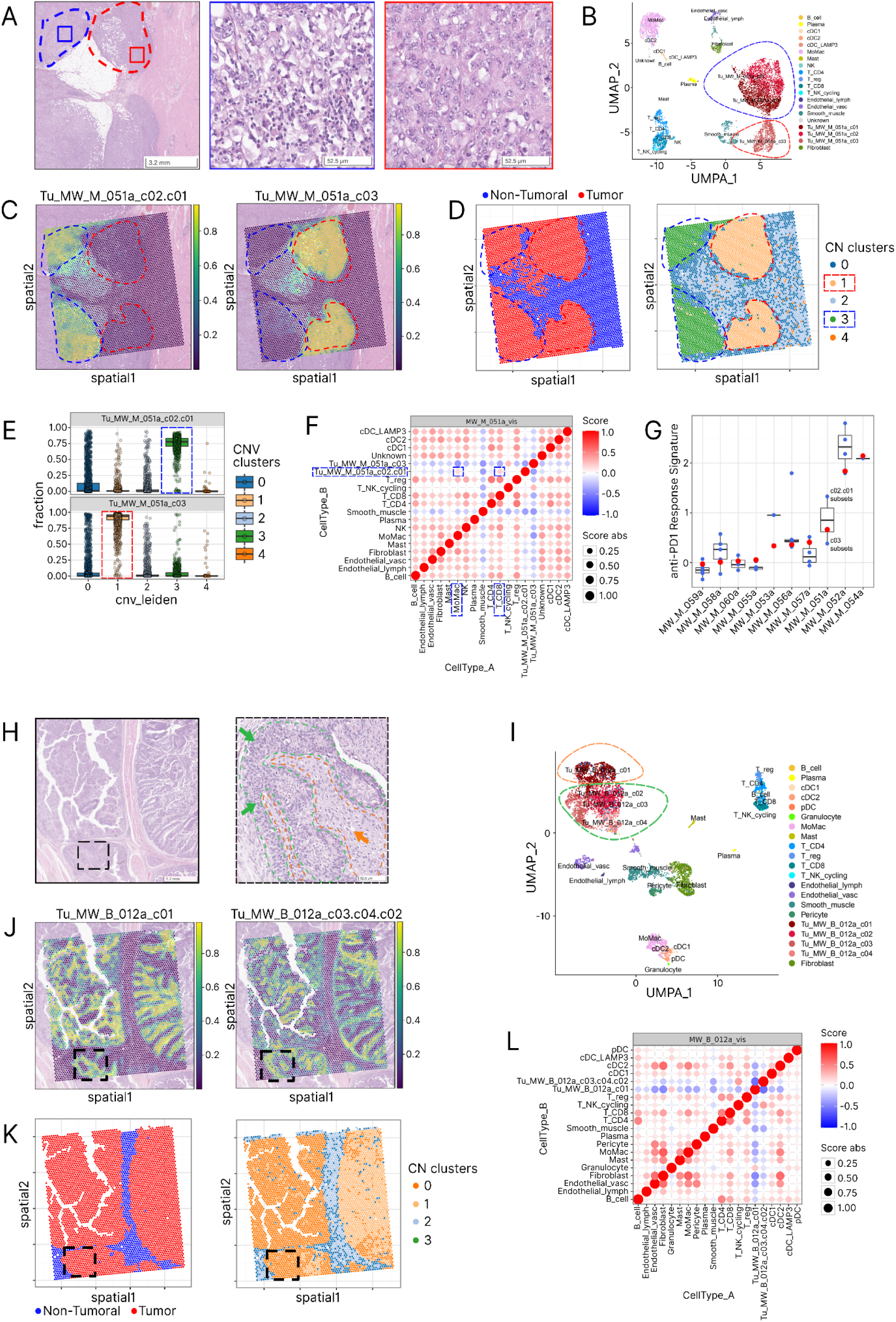
Multimodal integration of MOSAIC data highlighting the intra-tumor heterogeneity in mesothelioma and bladder tumor cases (Case #1 and Case #2) A. H&E image from a mesothelioma patient (MW_M_051) highlighting 2 different areas with red and blue. In the detailed view of the blue area (left), tumoral cells are similar to the red area, however a very prominent immune infiltrate is present. Detailed view of the red area (right), showing cells with abundant cytoplasm, large circular nuclei, and prominent nucleoli, without immune infiltrate. B. UMAP embedding from single cell experiment for sample MW_M_051a, cell color is based on the annotation of cell populations. Malignant cell cluster grouping is indicated by dotted line based on their colocalization (see Methods) C. Visium spot level deconvolution of malignant cell populations identified in single-cell for sample MW_M_051a. The two panels are corresponding to spatially distinct tumor cell populations, dotted lines corresponding to two identified areas in panel A. D. The left panel shows the spots used as reference for inferCNV in blue and observation in red, using a 10% tumor fraction threshold. On the right, the spots are colored based on the leiden clustering performed based on inferCNV results, displaying distinct regions based on their CN profiles. E. for the two patients, these boxplots indicate the distribution of each tumor cell fraction from deconvolution (see panel C) in each of the CN clusters (see panel D right). F. Bubble-plot showing the spot-wise Pearson correlation between each pair of cell populations in sample MW_M_051a. Point size corresponds to the absolute value of Pearson correlation while color is indicating the direction of the correlation. Bubbles and cell populations in dashed line squares are the colocalizations specific to one of the two subgroups. G. Boxplot indicating for the 10 mesothelioma patients of Mosaic Window, both the bulkRNAseq (red points) and the individual subpopulation in spatial transcriptomic (blue points) the activity of anti-PD1 response signature, using the signature established by R. Cristescu et al (Science 2018). For patient MW_M_051a, we used the Mann-Whitney statistical test to compare the distribution of signature activity between spots dominated by c01/c02 or c03 (p-value = 1.00:10-217). H. H&E image from a bladder cancer patient (MW_B_012a) focusing on a tumor region displaying an architectural pattern composed of two cellular subpopulations, arranged concentrically. The cells in the internal zone (orange arrow) exhibit cytological features as abundant eosinophilic/clear cytoplasm distinct from those in the peripheral area (green arrow) characterized by less cytoplasm and basophilic aspect, with pseudo vertical organization. Panels I, J, K and L are the same representations than panels B, C, D, and F, respectively, for sample MW_B_012a

SnRNAseq revealed three tumor clusters, which were assigned to two spatially distinct subsets based on co-localisation on ST, designated c01/c02 and c03 (Figure 3B-C). In addition, two spatially distinct copy number (CN) profiles were identified, mapping to the transcriptomic subsets and histological patterns (Figure 3D-E, Supplementary Figure 4).

Cell-colocalization analysis highlighted that the two regions are infiltrated by different non-malignant cell populations. The c01/c02 region is enriched with CD8+ T cells and macrophages, whereas the c03 region is relatively immune-depleted (Figure 3F, Supplementary Figure 5A). Coherently, differential expression and pathways enrichment analyses between these two transcriptomics subsets show enrichment in immune-related pathways associated with neutrophil degranulation, interferon and interleukin signalling in c01/c02 and extracellular matrix organization pathways in c03 (Supplementary Table G-H).

These findings suggest that CN profiles contribute to the tumor’s transcriptomic patterns and morphology.

We found a statistically significant higher activity of an anti-PD-1 response signature was detected in the c01/c02 cluster compared to c03. When comparing signature activity derived from bulkRNAseq with that of cancer subpopulations analyzed using ST, we observed significant intra-tumoral variability (Figure 3G, Supplementary Figure 5B-C). These findings underscore the potential of spatial transcriptomics to reveal mechanisms of treatment resistance and inform the selection of optimal treatment combinations. Furthermore, snRNAseq enrichment analysis between transcriptomic tumor subsets identified REACTOME SIGNALING_BY_MET pathway as a top enriched pathway in c01/c02 (p-value = 0.006, NES = 1.89, Supplementary Table H, Supplementary Figure 6A). These two observations highlight the presence of distinct tumor phenotypes and niches with potentially different sensitivities to immune checkpoint inhibitors and/or MET inhibitors. Interestingly, the enrichment of the MET signaling pathway in the c01/c02 subset aligns with the presence of neutrophil infiltration, which may contribute to MET activation via HGF secretion by neutrophil granulocytes. This mechanism has been implicated in fostering chronic inflammation, promoting tumor progression, and impairing response to immune checkpoint inhibitors, including anti-PD-1 therapy (Franciel Batista et al, 2023; Xia et al, 2025; Glodde et al, 2017). Our findings suggest that neutrophil-driven MET activation in this subset could underlie resistance to immunotherapy, reinforcing the role of the MET-HGF axis as a key player in the tumor microenvironment and a potential therapeutic target.

For c03 cluster, PROGENy pathways spot level enrichment analysis highlighted that this tumor region differentially activates pathways of extracellular matrix and has stronger inactivation of p53 pathway (Supplementary Figure 6C-D). The specific activity of MAPK and VEGF pathways is opening distinct opportunities for precision medicine in this c03 subset.

#### Case 2 - (MW_B_012a)

A 73-year-old male patient, a former smoker (15 pack-years), had a history of diabetes, hypertensive heart disease, hypertension, and stage III chronic kidney disease. He was overweight (BMI 29 kg/m²), had treated dyslipidemia, and mild hepatic steatosis. He was diagnosed with pT1 bladder cancer following a TURBT, without evidence of disease extension. Less than a week after diagnosis, he underwent an upfront cystectomy with lymph node dissection; which revealed a pT2N0 papillary invasive urothelial carcinoma without divergent/variant histology or lymphovascular invasion, and the resection margins were negative (R0). He died less than 2 months after diagnosis from an unknown cause. His most recent CT scan - done 22 days after diagnosis - showed no signs of disease recurrence.

The tumor exhibits a concentric architectural pattern with two distinct cellular subpopulations. The cells in the central region are composed of large cells with a geometric shape and well-defined borders. The cytoplasm is abundant and eosinophilic; moreover, many cells exhibit a perinuclear white halo. The nuclei are pleomorphic, hyperchromatic, and heterogeneous. On the other hand, the peripheral cells have less cytoplasm, appear more basophilic, and are arranged in a pseudo-vertical orientation, which resembles basaloid morphology (respectively orange and green regions annotated in Figure 3H).

SnRNAseq reveals four tumor transcriptomic clusters, combined to two subsets based on co-localisation on ST, designated c01 and c02/03/04 (respectively mapping the internal orange and external green regions; Figure 3H-J). From a CN perspective, we identified two distinct profiles corresponding to two large, spatially distinct cancer areas (Figure 3K, Supplementary Figure 7). Notably, the two major transcriptomic subsets observed by snRNAseq are present within each CN profile, and these transcriptomic profiles correlate with visibly distinct cytological patterns in both areas. These observations suggest that the transcriptomic profiles appear to drive the cytological phenotypes, while the CN profiles do not drive the transcriptomic states, suggesting that this different genomic landscape represents passenger alterations in this specific tumor.

TME analysis revealed that c01 cancer cells are co-localized with granulocytes (Figure 3L; Supplementary Figure 8) while c02/03/04 is overall cold without specific infiltration of immune cells.

Differential expression and pathways enrichment analyses between these two transcriptomics subsets showed an enrichment in keratinization and fatty acids metabolism for c01 subset (Supplementary Tables I,J), suggesting a probable squamous differentiation of the cells in this subset, which correlates with the previously described histological findings (Figure 3H). This observation aligns with studies linking keratinization pathways to basal-like urothelial carcinoma, which often exhibits squamous differentiation and an aggressive clinical course (Lan et al, 2015). c02/03/04 was characterized by an enrichment of cell cycle pathways suggesting more proliferative cancer cells in this region. The spot level PROGENy pathway enrichment analysis highlighted a significantly higher activity of hypoxia pathway in c01 subset, that could be associated with a possible maturation state towards squamous phenotype and also bad prognosis, while c02/03/04 was characterized by stronger inactivation of p53 and higher MAPK activity (Supplementary Figure 9).

#### Case 3 - (MW_B_012a)

A 59-year-old male patient in good general health—with a history of active alcohol consumption, former smoking (15 pack-years), and hypertension—was diagnosed with pT1 bladder cancer and grade 3 carcinoma in situ (pTis) following a TURBT, with no evidence of disease extension. Three months after diagnosis, he underwent an upfront cystectomy with lymph node dissection. Pathology revealed a pT3N1a solid invasive urothelial carcinoma, without divergent histology or lymphovascular invasion, and with negative resection margins (R0). He received three cycles of adjuvant platinum/gemcitabine chemotherapy; however, because the first cycle with cisplatin was complicated by acute renal failure, the subsequent two cycles were administered with carboplatin instead. Nine months after diagnosis, he developed locoregional and metastatic recurrence and was treated with palliative docetaxel. Two months later, he received a single injection of the anti-PD-L1 antibody atezolizumab for further disease progression, but this treatment was discontinued due to an allergic reaction. Following palliative locoregional radiotherapy, he died one year and six months after diagnosis from disease progression, compounded by renal and hepatic insufficiency.

This large nested bladder cancer exhibits a consistent histological pattern—tumor nests separated by stroma—with distinct cytological features. Within each nest, round cells with clear cytoplasm and minimal organization are observed, while at the periphery, cells are elongated and arranged in a vertical “palisadingˮ pattern. A characteristic feature of large nested carcinoma, central sterile comedo-like necrosis, is seen within the tumor nests. Additionally, tertiary lymphoid structures (TLS), which are often associated with this tumor type, can be observed between nests, with some appearing immature. Each nest is encased by a thick band of fibroblasts and immune cells (Supplementary Figure 10A).

snRNAseq identified four transcriptomic clusters, combined into two subsets based on spatial co-localisation on ST, designated c02/03 and c01/c04. These are distributed across two regions: the core and the edge of the tumor nests, respectively (Supplementary Figure 10B-C). In addition, a single CN profile was observed, localized to the core of the tumor islets (Supplementary Figure 10D and 11).

Cell co-localization analysis mainly confirmed the observed fibroblast layer surrounding the nests observed in the histopathological inspection (Supplementary Figure 10E and 12).

Differential expression and pathway enrichment analyses between these two transcriptomics clusters showed that c01/c04 subset is characterized by an activation of extracellular matrix organisation (adjusted p-value=0.008 NES=2,13, Supplementary Table K,L), consistent with the fibroblast layer co-localized with this cluster of malignant cell, and signaling by receptor tyrosine kinase (adjusted p-value=0.008 NES=2,08, Supplementary Table K,L) whose activation could be the consequence of Cancer Associated Fibroblasts activity toward cancer cells (Wu et al, 2021), in particular considering the leading edges of this enriched pathway, including MMP2, ITGA2, ITGA3 and ITGB4. On the other hand, subset c02/c03 is characterized by pathways related to innate immune system (adjusted p-value=0.008 NES=-2.22, Supplementary Table K,L) and neutrophil degranulation (adjusted p-value=0.008 NES=-1.92, Supplementary Table K,L) that could be explained by the presence of blood vessels in the core of the nests.

The histological pattern of ‘large nested’ is associated with FGFR3 activation in MIBC (Weyerer et al, 2020). To investigate potential genetic alterations, we examined bulk RNA sequencing data for gene fusions and whole-exome sequencing data for activating mutations. However, no FGFR3-related alterations were detected. Notably, FGFR3 expression was markedly elevated in this sample, suggesting activation through an alternative mechanism (Supplementary Figure 12).

The differential expression analysis also highlighted CEACAM5 and CEACAM6 as overexpressed by c02/c03 cell subset (Supplementary Table K), suggesting a potential for antibody drug conjugate (ADC) therapy targeting these molecules.

In summary, while the tumor presents a unique malignant cell subset from the copy number perspective, it displays divergent transcriptomic profiles corresponding to the cores versus the edges of the tumor islets and their surrounding cells.

#### Case 4 - (MW_O_016)

A 57-year-old postmenopausal female patient with no significant medical history other than a first-degree family history of colon cancer was diagnosed with FIGO stage IIIC epithelial ovarian high-grade serous carcinoma. She received three cycles of neoadjuvant carboplatin/paclitaxel chemotherapy, during which a partial response was noted, followed by interval debulking surgery via midline laparotomy, which resulted in no residual disease. Subsequently, she underwent three cycles of adjuvant chemotherapy with the same regimen, with the final cycle administered six months after diagnosis. Bevacizumab was initiated concurrently with the adjuvant chemotherapy and continued as maintenance therapy until one year and seven months after diagnosis. A disease recurrence was documented immediately after, for which she received eight cycles of palliative chemotherapy consisting of carboplatin/gemcitabine. She died of the disease three years after diagnosis.

Histologically, the tumor shows a single papillary pattern while the snRNAseq reveals two primary transcriptomic subsets designated c01 and c02, localised in different regions (Supplementary Figure 10F-H). The CN analysis indicated that only a single genomic profile exists in these two cancer regions (Supplementary Figure 10I and 13).

The cell-colocalization analysis suggests that the c02 subset is strongly associated with pericytes (Supplementary Figure 10J and 14) but this observation was not made in the histo-patholocal analysis.

Genes associated with activation of the pre-replicative complex and DNA replication are enriched in c01 and genes associated with cell cycle, mitosis, intracellular transport, membrane trafficking-related pathways are enriched in c02 (Supplementary Tables M,N). Furthermore, a statistically significant higher expression of an Homologous Recombination Deficiency (HRD) signature was detected in the c01 subset compared to c02 (Supplementary Figure 10K), highlighting the presence of distinct tumor niches with potentially different sensitivities to PARP inhibitors.

In summary, this patient showcases a unique cancer phenotype both at the histopathological and genomic level, whereas the transcriptomic analysis identifies two spatially distinct cancer cell populations, with a different predicted response to PARP inhibitors.

Overall, the multimodal investigation of these 4 cases clearly demonstrates the value of the MOSAIC dataset. We invite researchers to leverage this dataset to discover novel biomarkers, investigate mechanisms of resistance, and create predictive models for cancer therapies. Our goal with the MOSAIC Window Initiative is to foster collaboration and innovation in cancer research, ultimately leading to discoveries that improve patient care and clinical outcomes.

## Discussion

In this paper, we present the MOSAIC study and its framework, encompassing the initial consortium setup, data generation and integration processes, and the release of MOSAIC Window along with selected use cases demonstrating translational potential of multiomics data, with end goal to advance the application of AI for biological insights.

Thanks to the MOSAIC study we developed a unique, large-scale multiomics dataset designed to advance precision oncology by providing comprehensive pan-cancer molecular and spatial information.

This dataset represents a groundbreaking resource to address current limitations in ST data analysis, such as multimodal data integration and model efficiency (Durand, MIT Technology Review, 2023). While the biostatistics literature is increasingly rich in terms of methods for ST data analysis (see, e.g., the review of Zeng et al. 2022, and references therein), MOSAIC high-resolution approach through ST and snRNAseq enables more detailed studies of tumor microenvironment heterogeneity, allowing identification of rare cell populations and immune cell interactions that would be missed in bulk genomics data.

Even with smaller datasets ranging between 12 to 36 samples, this approach has proven valuable in various cancer studies (De Zuani et al, Nat Commun, 2024; Berglund et al, 2018; Andersson et al, 2021). For instance in prostate cancer research, ST allowed for better delineating of cancer foci which were not visible on the histological slides (Berglund et al, 2018). Similarly, in HER2+ breast cancer, integrating single-cell data with ST has led to discoveries about tertiary lymphoid structures and immune cell interactions, enabling the development of predictive models across different tissue types (Andersson et al, 2021).

Given its scale and ambitions, the MOSAIC project identified critical steps, as well as technical and operational challenges relevant for large multi-centre omics studies. Among these are selection of ST technology, based on criteria including full transcriptome capabilities, multi-tumor compatibility, and scalability (Dong at al., 2024), as well as management of operations between several sites both in the EU and US. From an organizational and governance point of view, an important insight was to recognize the importance of early involvement from all teams including clinical oncology, IT, data privacy organizations, and especially pathology, whose work is at the core of sample selection and data generation. Proactive engagement with IRBs and Ethics Committees is key to avoid regulatory delays, while adopting a patient-centric consent can boost participation rates.

In this work, we do not only provide a detailed overview of MOSAIC data generation and processing, but we also present the MOSAIC Window initiative, a sharing data which allows researchers to access 60 MOSAIC patients. In particular, we explore 4 cases from the MOSAIC Window dataset. The multimodal analysis presented has revealed intricate patterns of intra-tumor heterogeneity across mesothelioma, bladder, and ovarian cancers, each showing distinct molecular and cellular characteristics that could influence therapeutic strategies. The evidence demonstrates that integrated analysis of H&E slides, snRNAseq, ST, and copy number variations can unveil complex tumor architectures, as seen in Case 1 where distinct regions showed different immune infiltration patterns and potential therapeutic sensitivities. The findings are particularly compelling in cases like the bladder cancer specimens, where the analysis revealed relationships between genomic profiles and transcriptomic states. Moving forward, AI-driven approaches could automate the integration of these multiple data modalities across larger patient cohorts, potentially identifying recurring patterns of tumor organization and their clinical implications. Machine learning algorithms could be particularly valuable in recognizing common molecular signatures across different cancer types, helping to establish a systematic classification of tumor architectures. This automated approach would not only accelerate the analysis process but also potentially reveal new patterns that might be missed in case-by-case manual analysis, ultimately leading to more standardized and actionable therapeutic recommendations.

### Future directions for AI-enabled analysis of MOSAIC data

While this paper focuses on four examples to illustrate how deep molecular and clinical data can be integrated to extract novel clinical insights, the MOSAIC dataset holds far greater potential — particularly in light of recent advances in AI techniques.In particular, we see 4 areas where AI models could highly benefit from MOSAIC multimodal data.

#### Multimodal data integration

First, we aim at improving the usability and integration of ST and snRNAseq, to enable richer downstream analyses for researchers. The existing literature on analysis methods for snRNAseq is already rich and includes methods for processing the data (see, e.g., the literature review in Heumos et al. 2023), differential expression analysis (see Squair et al. and references therein), clustering methods (see Kiselev et al. 2019 and references therein), and recently foundation models pre-trained on large scale data sets (Yang et al. 2022, Zhang et al. 2022, Hao et al. 2024, Cui et al. 2024). Comparatively, existing methodologies to analyze ST are much less mature, given the novelty of the technology.

While leveraging the already available techniques for ST - covering relevant topics such as technical artefacts and batch effect removal (Ni et al. 2022, Giotto et al. 2024, Righelli et al. 2024), differential expression analysis (Svensson et al. 2018, Cable et al. 2024, Mason et al. 2024), dimensionality reduction (Shang and Zhou 2022, Li et al. 2023, Xu et al. 2023) and clustering (Dong and Zhang 2022, Sottosanti and Risso 2023, Long et al. 2023) - MOSAIC provides new opportunities to further explore multimodal integration.

When focusing on ST and snRNAseq, one interesting area of development is the application of “super-resolution methods,ˮ which use AI to increase the resolution of spatial omics assays to near single-cell levels (Zhao et al. 2021, Bergenstråhle et al. 2022).

More generally, data integration in biology has been often analyzed through standard multimodal approaches such as early, intermediate and late fusion (Boulahia et al. 2021), which boil down to concatenating the data before or after some modality-specific transformations, often lead to limited improvement compared to classical AI tools (Lin et al. 2023), while impairing the biological interpretability of models. On the contrary, successful multimodal AI approaches exploit prior knowledge to capture the complex biological interdependence between, e.g., RNAseq and histology data (Schmauch et al. 2020). Such methods usually handle multimodality in innovative ways; for instance, leveraging medical images to identify biologically meaningful regions within a spatial transcriptomics sample, before proceeding to spatialized differential expression analysis based on annotated regions (Cable et al. 2022).

In this sense, MOSAIC offers a unique resource to develop and test several approaches to multimodal integration, not only for ST and snRNAseq, but also for more diffused modalities such as H&E slides, WES and bRNAseq.

#### Spatially aware prognostic or predictive biomarkers of patients outcomes

A second promising stream that we believe will significantly benefit from the MOSAIC data is the development of spatially aware prognostic or predictive models.

Current research has identified significant gaps in two critical areas: the lack of efficient, interpretable predictive models for clinical outcomes using spatial omics, and the need for robust, large-scale evaluation of existing methodologies. By leveraging MOSAIC’s extensive scale, researchers can develop biologically meaningful spatially aware biomarkers to predict treatment responses and outcomes across large patient populations. This innovative approach could lead to breakthrough discoveries, such as identifying specific subpopulations with distinct anti-PD1 response signatures (Figure 3G). Such advancements would not only enhance our understanding of spatial biology but also potentially revolutionize personalized treatment strategies in clinical settings.

#### Target identification

Leveraging spatially aware methodologies on this multimodal dataset could highlight complex biology and enable the generation of relevant insights at the gene level, which are highly valuable for prioritizing therapeutic targets. For example spatially aware differential expression analysis could highlight genes having an important role in tumor progression (D.M. Cable et al. 2022; V. Svensson et al. 2018). By leveraging MOSAIC’s spatial and molecular insights, researchers can better understand and characterize intra and inter-tumor heterogeneity, ultimately leading to the identification of relevant biological mechanisms that can be targeted with precise therapeutic strategies. Furthermore, spatial transcriptomics enables a deeper understanding of the tumor microenvironment and tumor evolution, facilitating the identification of effective combination treatments tailored to specific tumor contexts. By capturing these dynamic interactions, this dataset has the potential to facilitate target identification and drive advancements in personalized oncology.

#### Foundational models

Last but not least, another encouraging approach towards multimodal AI is the development of foundational models for biology, for which MOSAIC data offers an unprecedented training opportunity. Indeed, recent advances have demonstrated the potential of foundational models for histology images (Filiot et al. 2024), single-cell transcriptomics (Cui et al. 2024), but also multimodal data including spatial transcriptomics (Lin et al. 2024). Thus, training such foundational AI models on the extensive data from the MOSAIC cohort, where both spatial gene expression profiles and corresponding histology images are available, is a very promising direction. By learning the intricate relationships between the visual features in H&E images and the underlying spatial transcriptomics data, AI models can be trained to, among other things, accurately reconstruct gene expression profiles from these widely available histological images (Schmauch et al. 2024). After validation, these models can analyze millions of H&E images routinely collected. This would expand spatial transcriptomics data availability without expensive experimental procedures. Through larger sample sizes, AI could deepen our understanding of tumor biology by revealing patterns invisible in smaller datasets. This approach could accelerate biomarker and therapeutic target discovery by creating a comprehensive virtual dataset spanning diverse cancer populations and stages.

## Methods

### Methods A: MOSAIC consortium

#### A.1. Sample qualification and lab workflow

As specified in the “Cohort mappingˮ section, a patient can be included in MOSAIC only when fitting into one of the approved cohorts. Once a cohort is approved (i.e., through ‘Cohort Selection’), the detailed identification of samples begins at the FP under ‘Sample Inclusion.’ The process starts with generating a patient list by screening Electronic Health Records (EHR) for inclusion and exclusion criteria, verifying consent eligibility, and confirming sample availability at the Biological Resource Center (BRC). Next, FFPE blocks, along with the corresponding archival H&E slides, are sent to the pathology laboratory to ensure that the samples meet the required specifications. Briefly, the steps are the following: I) Collect all available tumor blocks and (digitized) H&E for best block selection; II) H&E review by the pathologist and selection of the optimal block; III) Macroscopic quality check of the selected block and inclusion in the eCRF to create the unique patient and sample IDs; iv) Marking of the 6.5 x 6.5 mm area for the Visium frame on the H&E slide/image, and, if needed, the area of the slide to guide macrodissection. If archival H&E slides are unavailable, hematoxylin-eosin-safran (HES) slides can be used as an alternative for block selection. In specific cases, specimens that do not meet the minimum 40% tumor burden required by the inclusion criteria are re-embedded to ensure a consistent tumor percentage across all techniques.

The most complex step is the sample review as tissue requirements differ between solid tumors and DLBCL, in addition to some specificities associated with the sample types [biopsies; transurethral bladder resections (TURBT); surgical specimens]. The majority of the samples were under 10 years old at inclusion; had a minimal depth of 125 µm and minimal surface covering at least 20% of the Visium frame, with a preference for a minimum 25 µm². Cytologies/cytoblocks that do not preserve the tissue architecture were excluded. The common criteria for solid cancers is a tumor burden between 40 and 80%. The corresponding criteria for DLBCL is a minimum of 80% high grade component. For DLBCL samples not harvested from lymph nodes, macrodissection is applied to minimize contamination by healthy adjacent tissue. Cases with intermingled tumor cells and the healthy organ specific tissue were excluded.

It is important to note that assessment of these criteria was done in a two step manner, first in the 6.5 x 6.5 mm area selected for the Visium frame, relevant for ST; second on the whole slide (unless the sample was entirely included in the Visium frame area), relevant for bRNAseq, snRNAseq, and WES. The need for macrodissection was adapted accordingly (Supplementary Figure 3B-C).

Once enough blocks were identified for a given cohort, they were distributed into weekly batches of 8 to 16 samples to enter the data generation workflow, a step defined as ‘Data Generation Plan’. The clinical data associated with these samples was used to balance the batches, such as survival outcomes, histological or molecular subtypes, timepoints of harvest (e.g., baseline vs recurrence), to prevent any batch from being linked to specific criteria that could impede confident batch correction in subsequent analyses.

In addition to the challenge of creating a common eCRF that could accommodate both known and yet-to-be-defined tumor types, we also faced center-specific challenges, such as: I) the heterogeneity in the availability of easily accessible information across different centers for identifying cohorts, obtaining associated samples, and preparing data generation plans; II) the need for consent check or first consenting; III) the need for block retrieval for older samples stored remotely and received by batches (not always in the expected order); IV) the lack of space in the labs for block storage given the MOSAIC project’s high-throughput. As a mitigation strategy, we collectively adapted the weekly data generation plans to each FP specificities.

Upon receipt of the FFPE tissue samples in the lab, each sample is assigned a unique identifier through the eCRF and entered into the Sample Tracking System (STS) to ensure accurate tracking throughout the workflow. No sensitive data is collected within this system, only metadata to describe the experimental conditions in which the samples are processed.

The STS is utilized to monitor and document every stage of the process, from initial receipt to final shipment of materials to the CRO for sequencing. All procedures, including the preparation of tissue sections, the Visium workflow, and the completion of the STS records, are described and standardized in the SOPs.

The FFPE blocks are first subjected to sectioning, where six consecutive sections are made. The workflow for these sections is as follows:

I. The first three sections of the block (4 µm thick for solid tissues, 3 µm for DLBCL) after trimming were prepared and stored as back-up sections for potential future use in target validation, or used for replacing failed H&E or Visium assays.
II. An extra H&E section (4 µm thick for solid tissues, 3 µm for DLBCL) was made to serve as the main section for H&E staining and for alignment during Visium data analysis. The staining was performed according to the routinely used local protocol.
III. A Visium section (5 µm thick for solid tissues, 4 µm for DLBCL) designated as the primary section for the Visium Spatial Gene Expression workflow.
IV. A Visium back-up section (5 µm thick for solid tissues, 4 µm for DLBCL) was to be used if the main Visium section failed during the workflow.

During the project, the sequence of sectioning was modified: the three back-up sections were followed by the extra H&E section, then the Visium section, and lastly, the Visium back-up section. This change was implemented based on analyses showing no statistically significant impact on image alignment quality. This modification streamlined the process, reducing the time required for sectioning and minimizing material loss.

Following the preparation of sections, curls are cut from the remaining FFPE block in the following order:

I. Two curls for snRNAseq analysis.
II. Two curls for bRNAseq analysis.
III. Two curls for WES analysis.

In the European sites, the two curls for bRNAseq and the two curls for WES are placed in the same tube, since the CRO workflow has one isolation step for both bRNAseq and WES.

The Visium sections are processed up to either the pre-amplification step in the European centers or the final library preparation stage at the University of Pittsburgh. Chromium workflow in the US is performed until the library preparation. Subsequently, the libraries are shipped to the CRO along with the curls designated for bRNAseq and WES for further processing and sequencing (Supplementary Figure 15).

The main stages of the Visium throughput include low throughput of 8 samples per week with two samples processed per run per operator, twice weekly, and upscaled throughput of 16 samples per week by increasing the number of samples processed per run to four per operator.

The STS is integral to managing all aspects of sample processing across the centers. It provides comprehensive tracking of each sample through the following steps:

I. Receipt of FFPE Block: The initial receipt and registration of each block into the STS.
II. Sectioning: Documentation of sectioning details (section thickness), including the date, section sequence, and if macrodissection/re-embedding was needed.
III. Visium Processing: Detailed tracking of Visium protocol steps, including the date of each step, operator information, lot numbers of reagents used, operator name and any deviations from standard procedures, along with corrective actions taken.
IV. Final Shipment: Coordination and logging of the final shipment of materials, including Visium libraries, Chromium libraries and curls for bRNAseq and WES, to the CRO for sequencing.

STS filling instructions are part of the SOPs that describe the manipulations in the lab. The STS enables continuous monitoring of sample processing efficiency and throughput. Regular audits of the STS records ensure compliance with project protocols and address any deviations promptly. This approach ensures that the high-volume processing of samples is conducted efficiently and accurately, with optimal data quality across the collaborating centers.

In addition to the user-facing interface, the STS also includes an internal-facing interface that serves as a project management tool. Specifically, the STS is integrated with our internal data platform, *Owkin K*, allowing us to track the data lifecycle, especially timelines, from the shipment of physical samples to the CRO, through to data reception on Owkin K. This integration ensures that no physical material is overlooked or wasted, particularly in time-sensitive matters. Additionally, data captured by the STS is also extracted and imported in Owkin K to be integrated into the data analysis to correlate output quality with specific lab processing factors.

#### A.2. Data generation details and references for each data modality

MOSAIC includes the following modalities:

I. Clinical data
II. Hematoxylin and Eosin (H&E) microscopic images
III. Spatial transcriptomics (ST)
IV. Single-Nuclei RNA transcriptomics (snRNAseq)
V. Bulk Ribonucleic Acid Sequencing (bRNAseq)
VI. Bulk Whole Exome Sequencing (WES)

I. **Clinical data:** The clinical data include detailed information such as demographics, medical history, cancer and treatment-related information, date and nature of sampling, and oncologic events during follow-up with an optimized collection of cancer outcomes. Data is collected via an electronic Case Report Form (eCRF) at inclusion in MOSAIC, capturing all relevant history between date of cancer diagnosis and inclusion date. For patients who are alive and under follow-up at the time of inclusion, clinical data—particularly oncologic outcomes—are collected annually for five years after inclusion, within a window of ±30 days from the inclusion date. Additionally, data are collected at the end of the study, which occurs at a maximum of six years post-inclusion, on the date of death, or if the patient is lost to follow-up, whichever comes first. The common forms for patient-related information are: I) Demographics (date of diagnosis, date of last follow-up or death, general demographic information, and the tumor subtype); II) Consent & Eligibility; III) Subject history; IV) Family History; V) Treatment form (capturing all possible oncological treatment types, dosage, routes, dates and response); VI) Oncologic events before inclusion in MOSAIC (progression/recurrence, other cancer); VII) Concurrent treatment form; VIII) ‘End of study’ form (capturing the cause of end of study or death, if applicable); IX) ‘Biobanked data and specimens beyond MOSAIC’ form (collects information on other available biobanked specimens in FPs for each patient, independently of MOSAIC); and, for patients still alive at the inclusion in MOSAIC, X) Follow-up form, collecting yearly the occurrence of novel oncologic events and/or death. The tumor type-specific forms are: I) Clinical (height, weight, date of diagnosis, tumor and metastasis location, cTNM, stage); II) Biology (blood +/- urine test results); III) Extra Tumor Treatment information; IV) Pathology (histological type & subtype, (y)pTNM, prognostic histological features, +/- immunohistochemistry (IHC) and fluorescence in situ hybridization (FISH) results); V) Mutations (capturing all known genetic alterations). The TNM stages appear as reported in the patients’ medical record and are from the 7th AJCC edition, and, for cases with samples from after 2016, from the 7th or the 8th AJCC edition.
II. **HGE**: The imaging data are H&E microscopic images, generated with Leica Aperio CS2 scanner with 40X magnification in all the participating centers to minimize batch effects due to different scanners. Standard site-specific staining protocols are used to reflect the variation across pathology institutes and allow models to adjust for out-of-domain effect. Complementing H&E images are captured with routine diagnosis H&E protocols on the Visium slides to aid data integration. Pathologist annotations on the tissue structures are also collected.
III. **ST**: The core modality of the MOSAIC Study is ST through the Visium Cytassist standard definition protocol from 10X Genomics. Visium Spatial Gene Expression is a next-generation molecular profiling solution for classifying tissue based on hybridization of gene-specific probes of over 18,000 genes in human samples. With a total capture area of 6.5 x 6.5 mm, Visium defines 5000 spots with a 55 µm resolution in diameter corresponding to about 1-10 cells on average (Ref: LIT000128 - Rev C - Product Sheet - Spatial biology without limits: Spatially resolve gene expression in FFPE samples). The 6.5 x 6.5 mm area is selected by pathologists according to the following criteria, ordered by priority: good representation of the tumor with a minimal tumor burden of 40% and covering at least 20% of the selected area; adjacent non-tumoral tissue if possible, invasive margin if possible, TLS if possible. The method was set up in five participating hospitals with manufacturer’s approved protocols. Up to eight slides are processed in parallel. Samples are sent to a contract research organization (CRO) after the pre-amplification step in Europe (CeGaT) and after library preparation in the US (GENEWIZ by Azenta Life Sciences). For all the samples, 22 PCR cycles are used in the library preparation with no qPCR step. Samples were sequenced at 25,000 reads per spot covered by tissue. If sequencing saturation fell below 50%, samples were sequenced once again with the same parameters.
IV. **snRNAseq:** Since the Visium platform used is not single-cell resolution, spatial data is complemented with transcriptomes from nuclei isolated from the FFPE blocks. This snRNAseq adapted from the Chromium FLEX protocol from 10X Genomics for FFPE tissues uses similar probe-based technology as Visium (Vallejo, A. F. et al., 2022). Curls (2 x 25 µm) are shipped to CeGaT in Europe, and processed locally in the US. Manufacturer’s protocols are used for dissociation with a gentleMACS Octo Dissociator. Nuclei are counted with Cellexa in Europe and Luna FX7 in the US. Pools of 4 or 16 samples were processed in one well, aiming for the recovery of 10,000 or 8,000 nuclei correspondingly in each sample. Samples are sequenced at CeGaT and GENEWIZ with 10,000 reads per cell.
V. **bRNAseq:** Bulk RNA sequencing is performed to assess the general transcriptome of the tumor, and to be able to compare the cell-level data with earlier studies that employed only bRNAseq methodology. 2 x 10 µm curls of FFPE tissue are shipped to the core facilities followed by RNA isolation (for EU sites a total of 40 µm is shipped in the same tube to perform both bRNAseq and WES). At CeGaT, RNA extraction is performed using the RNeasy FFPE Kit (Qiagen), following the manufacturer’s protocol. The MagMax FFPE DNA/RNA Ultra Kit (Applied Biosystems™) is used by GENEWIZ for the sequential isolation of both DNA and RNA from the same FFPE tissue samples. Post-extraction, RNA libraries are prepared using the SMART-Seq Stranded Kit (Takara Bio). The prepared libraries are then sequenced on the NovaSeq 6000 platform at CeGaT and NovaSeq X platform at GENEWIZ, using a paired-end read format with a read length of 2 × 100 bp with NovaSeq 6000 and 2 x 150 bp with Novaseq X, to achieve a targeted output of 50 million clusters per sample.
VI. **WES:** WES targets the protein-coding regions, known as exons, of an individual’s genome. By only sequencing the exome, which constitutes approximately 1-2% of the genome but harbors most disease-causing mutations, WES efficiently identifies genetic variations linked to disease. As with the bRNAseq, 2:10µm FFPE tissue curls are used for DNA isolation at CROs (for EU sites a total of 40 µm is shipped in the same tube to perform both bRNAseq and WES). At CeGaT, DNA is extracted using the QIAamp DNA FFPE Tissue Kit (Qiagen). In contrast, GENEWIZ utilizes the MagMax FFPE DNA/RNA Ultra Kit (Applied Biosystems™) for sequential DNA and RNA isolation from the same FFPE tissue samples, requiring between 2 x 10 µm and 4 x 10 µm curls (two for RNA and two for DNA extraction). For the library preparation, the Twist Comprehensive Exome Panel is employed, which targets exonic regions for sequencing. In addition to the exome panel, CeGaT also utilized the Twist Mitochondrial Panel to target mitochondrial DNA regions. Sequencing is carried out on the NovaSeq 6000 platform in CeGaT and NovaSeq X platform at GENEWIZ with paired-end reads of 2 × 100 bp and 2 x 150 bp correspondingly. The sequencing is configured to produce a targeted output of 100 million mapped reads per sample.

#### A.3. Data flow, processing and quality control pipelines for each data modality

One of the key advantages of the MOSAIC study is the centralization of all the data collected and generated in Owkin’s AI and data platform, *Owkin K*. Its purpose is to store, access and select MOSAIC data and host MOSAIC computational projects execution.

All data types integrated in the central platform are linked using common Subject IDs and Block IDs to allow accurate analysis and interoperability in accordance with FAIR principles (Findable, Accessible, Interoperable, Reusable). Metadata collection from the lab processes through the MOSAIC sample tracking system (STS) is also automatically exported to Owkin K to allow evaluation of biases and batch effects.

The architecture of the data platform is documented in the MOSAIC Data Management Plan and is represented in Supplementary Figure 1. The Owkin K platform supports the confidentiality, integrity and availability requirements for the study data as well as providing analytical capabilities and the ability to store derived data alongside the original data. The system is designed to ingest, pre-process, and store multi-modal data with different formats at scale, presenting integrated subsets of data to accredited and authorized researchers. Owkin K enables collaborative projects between FPs and Owkin or between two or more FPs, allowing a secure and shared analytical environment for scientific collaboration on data from multiple centers.

All MOSAIC modalities are generated through center sample preparation and CRO sequencing resulting in the central ingestion of raw data on the Owkin K data platform. These raw data include Images (H&E and ST modalities) and fastq files (WES, RNAseq, snRNAseq, and ST modalities).

For each sample block, data resulting from one modality is processed separately through a dedicated processing workflow relying on state of the art methods as detailed below.

I. **Clinical data** Clinical data is collected through a dedicated eCRF, which provides automatic checks on the fields present in each patient’s form (for example consistency checks, format checks, drop down menus). The data collected through the eCRF is automatically extracted and ingested in the central MOSAIC platform, *Owkin K*, on a daily basis. Once ingested, the data is processed through an automated pipeline which calculates some additional metrics, such as Overall Survival (OS) and Progression Free Survival (PFS). The first QC round combines automated checks and oncology expert review to identify inconsistencies and missing information. For instance, if “Death related to cancer” is marked as true without any recorded progression or recurrence before death, the system flags this missing progression event for human review. In such cases, Owkin contacts the center, the patient medical record is reviewed, the eCRF is corrected, and the data updated on Owkin K undergoes a second round of QC. All criteria deviations are tracked in a specific dashboard, which includes a description of the deviation, the final decision after discussion with the FP, and the updated result after review.
II. **HGE and ST:** For imaging data, we start by ensuring basic coherence in the set of images: the capture zone is located by the pathologist on the qualification H&E slide and the corresponding area in the slide adjacent to the Visium area, stained using regular H&E stains, is not degraded. On top of that, this adjacent H&E slide is semi manually checked (trained models are used to flag issues such as microscope focus failure or presence of artifacts such as penmarks, folds or tears), then human operators review each slide using these insights. For ST, an internal pipeline based on 10x SpaceRanger count v3.1.0 is used to process data, including the alignment between transcriptomics and the high resolution H&E image. Finally, both sample level and spot level QC thresholds are applied in order to remove low quality data. At the level of the sample, we ensure at least 50% of sequencing saturation, a median of at least 10,000 reads per spot, a median of at least 500 genes per spot and less than 20% of UMI estimated to come from genomic DNA. At the spot level, we filter out spots with less than 200 UMI to ensure high signal over noise ratio.
III. **snRNAseq:** 10x Genomics Cell Ranger v7.1.0 was used to demultiplex reads and generate gene expression matrices for each sample. Samples with low quality are removed (less than 500 cells or less than 50% of reads mapping to the probeset).
IV. **bRNAseq:** We use Owkin’s RNAseq pipeline that is based on best practices. RNAseq reads are aligned on human reference GRCh38 using STAR v2.7.8a. At this level, samples with more than 40% ribosomal RNA are excluded for downstream applications. Raw read matrix counts are then generated taking into account strandedness of libraries using gene positions and strands from Gencode databases (GRCh38_gencode_v37). Normalisations are then processed from this matrix including FPKM, TPM and DESeq2 VST (Love et al, 2014). To call gene fusions, we are using STAR outputs combined with STAR-Fusion v1.13.0 with default parameters.
V. **WES:** First, we use the Illumina DRAGEN Somatic Pipeline v4.2 in tumor-only mode to perform read alignment on human reference GRCh38. At this step, we exclude samples with low coverage of the capture exome regions, using 50X as minimal depth of sequencing. SNP/INDEL variant calling is also performed by the DRAGEN workflow and annotated with the Ensembl Variant Effect Predictor (VEP). In order to enrich calling in somatic alterations, common variants in germline databases gnomAD V4 are flagged as germline, using a frequency threshold of 1:10^-4^ in any of the populations defined in GnomAD database. We further enrich for oncogenic and likely-oncogenic mutations following heuristics based on the affected gene’s role in cancer, mutation effect on the protein and recurrence in large datasets of tumor samples. In short, genes affected by potentially oncogenic alterations are identified in each sample. A gene is said to be affected by a potentially oncogenic alteration in a given sample if it satisfies at least one of the following criteria:
  - The variant introduces a nonsense change in a known tumor suppressor gene (TSG). TSGs are obtained from IntOGen (Gonzalez-Perez, 2013). Nonsense changes are defined as variants with a high impact consequence from VEP annotations.
  - The variant produces a previously reported amino acid change in known cancer hotspots. Hotspots and reported amino acid changes were obtained from cancerhotspots.org and Hess et al. (Hess et al, 2019). Only SNVs in protein-coding genes were considered when analyzing hotspots.
VI. If a variant is flagged as germline with a frequency below 0.01 in any GnomAD population, but is potentially oncogenic following abovementioned heuristics, then it is whitelisted and included in the final VCF file.
VII. For CNV calling, we also use the Illumina DRAGEN CNV pipeline outputs to define samples with either a gain (more than 2 copies) of oncogenes or a loss (less than 2 copies) of TSGs, using the same resources as for variant calling.

## Methods B: MOSAIC Window

### B.1 MOSAIC Window patients description

Bladder: All 15 BLCA patients had samples obtained from an upfront cystectomy for stage II to IIIB disease. All but one patient underwent complete lymph node dissection. Only one patient had squamous cell carcinoma; the remainder had urothelial carcinoma, with three patients showing squamous differentiation. Additionally, one patient presented with a micropapillary variant and another with a sarcomatoid variant. Following surgery, three patients received adjuvant platinum/gemcitabine chemotherapy, and four were treated with immune-checkpoint inhibitors (ICI) upon relapse.

Ovarian: Among the 15 patients with ovarian serous high-grade carcinoma (stage IIB to IVA), including two with a known germline BRCA2 mutation, two samples were obtained from a post-NACT primary tumor, six from a relapse lesion, and the remainder from baseline surgeries or biopsies. Twelve patients received Bevacizumab as first-line maintenance therapy, while one patient was treated with a PARP inhibitor. Three patients experienced a platinum-resistant relapse, and three received ICI either as part of first-line maintenance or at relapse.

Glioblastoma: All 10 GBM samples were obtained from baseline surgeries, with primary tumor diameters ranging from 27 to 80 mm. Four patients had a methylated MGMT status, and all were IDH wild-type. All but one patient received adjuvant chemoradiotherapy with Temozolomide. One patient was treated with adjuvant Bevacizumab, while five others received Bevacizumab at relapse.

DLBCL: All 10 DLBCL samples were obtained from a baseline biopsy or surgery before treatment with first-line R-CHOP (Rituximab, Cyclophosphamide, Doxorubicin, Vincristine, and Prednisolone) chemotherapy for Ann Arbor stage III or IV disease. Baseline LDH levels ranged from 197 to 980 U/L. Two patients had the activated B-cell-like (ABC) subtype, six had the germinal center B-cell-like (GCB) subtype, and the subtype was unknown for two patients. At relapse, one patient was treated with CAR-T cell therapy.

Meso: All MESO samples, except one, were obtained from baseline biopsies or surgeries for stage I-IV diffuse mesothelioma; the remaining sample was collected from a post-NACT surgical specimen. In total, four patients received NACT, resulting in one partial response and three instances of disease progression. Additionally, half of the tumors were epithelioid while the other half were of the biphasic subtype, with nuclear grades ranging from I to III. At relapse, five patients were treated with ICIs.

### B.2 MOSAIC Window data processing for the 4 use case analysis

In order to fully leverage the multimodal nature of MOSAIC data (see Methods A for data generation and processing details), we followed a five steps approach, analyzing one modality on top of the other and then capturing the TME complexity and variability through integration of insights coming from the different data layers.

#### Histopathological evaluation

First, a systematic histopathological review of the H&E stained sample was conducted by an expert pathologist, following the standards established by the World Health Organization (WHO) for tumor classification. This evaluation assessed tumor heterogeneity in the tissue. In cases where multiple tumor regions were identified, detailed segmentation was performed through precise histological annotations, documenting predominant cytological patterns in each tumor region, degree and distribution of immune and stromal cell infiltration, as well as cellular and structural organization, including tissue architecture, neoplastic cell arrangement, and the presence of morphologically relevant features. These annotations were used as a reference in the following analysis, serving as a baseline for comparison and validation of results obtained through computational and experimental approaches. This step established a histopathological quality control framework, ensuring data integrity and interpretations validity.

#### snRNAseq processing and analysis

For the analysis of the four cases, we used the processed 10x Genomics Cell Ranger outputs after cell and genes filtration and counts normalization (Methods A.3.III.).

From these count matrices, SoupX v1.5.2 (Young et al, 2020) was used to estimate the ambient RNA expression profile from empty droplets and remove cell free RNA contamination in the data, considering 50 principal components. Cells with mitochondrial, ribosomal or hemoglobin genes content over 10%, fewer than 200 genes expressed and more than 6000 genes expressed were removed in the QC process. Following cell filtration, we removed genes that were expressed in fewer than three cells across the entire sample to eliminate potential technical artifacts. We then used scDblFinder v1.4.0 (Germain et al, 2021) to flag and remove multiplet cells using the default number of 19 principal components. Counts were then normalized using SCTransform implemented within Seurat v4.1.1 (Hao et al, 2024).

Finally, individual sample objects are merged at the level of the pool and dataset to perform cell type annotation by using scVI (Lopez et al, 2018) using a reference dataset manually annotated based on known biomarkers (Supplementary Table E) at high granularity (Level 4, Supplementary Table F) all indications except GBM. For GBM, the GBmap atlas (Ruiz-Moreno et al, 2022) was used as a reference dataset. The cell type annotation Levels 1-3 were added according to the hierarchy rules (Supplementary Table F). For cancer cell populations, a shared nearest neighbour (SNN) modularity optimization based algorithm with a resolution of 0.4 was used to automatically identify transcriptionally distinct populations within each donor. Any cancer cell clustering inside a non tumor cluster was flagged not otherwise specified (NOS) and dropped from downstream applications. In the four cases selected for this paper, an average of 4.5 clusters of tumor cells were identified with this approach. Finally, differential expression analysis (DEA) between these sub-populations and minimization of the condition number was conducted to generate signature matrices using Seurat FindMarkers function with default parameters and the kappa function. Values reported in the signature matrix correspond to log2 fold change of significantly differentially expressed genes (adjusted p-value < 0.05) and with a positive log2 fold change.

#### Spatial transcriptomic processing and integration with snRNAseq

For the analysis presented in this study, we used the processed 10x Genomics Space Ranger outputs of MOSAIC Window dataset (Methods A.3.II.). Scanpy v1.10.1 package was used to perform CPM and logCPM normalisations, respectively using scanpy.pp.normalize_total (target_sum=1,000,000) and scanpy.pp.log1p methods.

To spatially map cell population at the transcriptomic level, we used cell2location v0.1.3 (Kleshchevnikov et al, 2022) deconvolution method on raw counts and the indication specific signature matrices generated in the snRNAseq processing (c2l.models.Cell2location: N_cells_per_location=8; model.train: max_epochs=10_000 and train_size=1; model.export_posterior: num_samples=2000 and batch_size fixed number of spots in the Visium sample). In order to identify spatially distinct cancer cell populations for each sample, we iteratively regrouped those with a spotwise Pearson correlation above 0.5, summing the cell2location weights of aggregated cell population. Cell type weights at the spot level were normalized to obtain cell fractions for each Visium spot.

#### Transcriptomic characterization of cancer sub populations

After identification of spatially distinct tumor cell population, we used snRNAseq data to perform DEA using Seurat’s FindMarker with default parameters by comparing each population with all the others. Genes with a significant (threshold adjusted p-value < 0.05) were ordered on the log2 FoldChange to perform pathway enrichment analysis using fgsea v1.16.0 (Sergushichev, 2016) with REACTOME gene sets (Milacic et al, 2014).

To characterize tumor sub-population specific TME, we calculated the pairwise Pearson correlation for each pair of cell populations using calculated cell fractions.

#### Identification of genomic subclones and mapping with transcriptomic phenotypes

To characterize tumor regions at the genomic level, We used infercnvpy v.0.6.0, a python implementation of the original inferCNV method (Slyper et al, 2020), to discover potential genomic subclones by identifying distinct copy number (CN) profiles of malignant cell, which are then mapped into their spatial distribution. Spots with more than 500 UMIs and <10% Tumor fraction at level_2 deconvolution (Supplementary Table F) was used as reference. Gencode v47 transcriptomic reference with 500 genes bins were used for the modelisation of copy number profiles. Leiden clustering (resolution=0.2) of CN profiles was performed to aggregate spots into similar profiles allowing the identification of distinct tumor sub-clones within each patient. To identify potential mapping between genomic and transcriptomic regions, we studied the distribution of the different transcriptomic cancer cell fractions in theCNV clusters.

#### Quantification of pathway or gene signature activity

In order to better understand the specific biology of each spatially distinct population we performed spot level quantification of gene sets and gene signatures of interest.

Using decoupler v1.8.0 (Badia-i-Mompel et al, 2022) get_progeny (top=500 genes representing each pathway) and run_mlm modules, we performed spot level activity of all pathways contained in PROGENy database (Schubert et al, 2018).

In addition, we used two gene signatures from literature: Homologous Recombination Deficiency (HRD) signature from G. Pang et al. (Peng et al, 2014) and anti-PD1 response signature from R. Cristescu et al. (Cristescu et al, 2018). Spot level quantification of these two signatures was performed using run_ulm method from the decoupler package.

In order to measure the differential activity of these gene sets or gene signatures in any of the spatially distinct transcriptomic cancer populations, spots with a dominant fraction in any of the cancer populations were compared. Mann-Whiteney statistical test was used to measure the significance of these differences.

To compare the activity of the anti-PD1 signature between bulkRNAseq and spatially aware subpopulations, we analyzed data from 10 mesothelioma patients in MOSAIC Window. We assessed the signature activity defined by R. Cristescu et al. at both the bulk level and within cancer subpopulations. Subpopulations were identified using the same approach as for the four cases (see Methods: Transcriptomic Characterization of Cancer Subpopulations). We then calculated the signature activity for bulkRNAseq and for the average of spots dominated by each subpopulation using the run_ulm method from the decoupler package.

#### Multimodal integration

All the insights generated in the previous methodology were co-analyzed and integrated to achieve a multimodal view of the MOSAIC Window samples to showcase the value of MOSAIC multimodal data for characterizing patient intra-tumoral heterogeneity and identifying cellular niches and their impact on communication mechanisms determining cancer development and resistance. We utilized multiple complementary approaches to analyze tumor heterogeneity: HE for microscopically different tumor regions and cytological aspects, snRNAseq for malignant cell subclusters, and ST to map - and possibly merge - these subclusters spatially, defining the final number of malignant cell transcriptomic phenotypes. Using inferCNV on ST data revealed genomic clone distribution. By integrating these analyses, we examined the relationships between histological, transcriptomic, and genomic features, while also studying differential gene expression and pathways in tumor regions and their microenvironment.

## Supplementary Figures

**Supplementary Figure 1.**
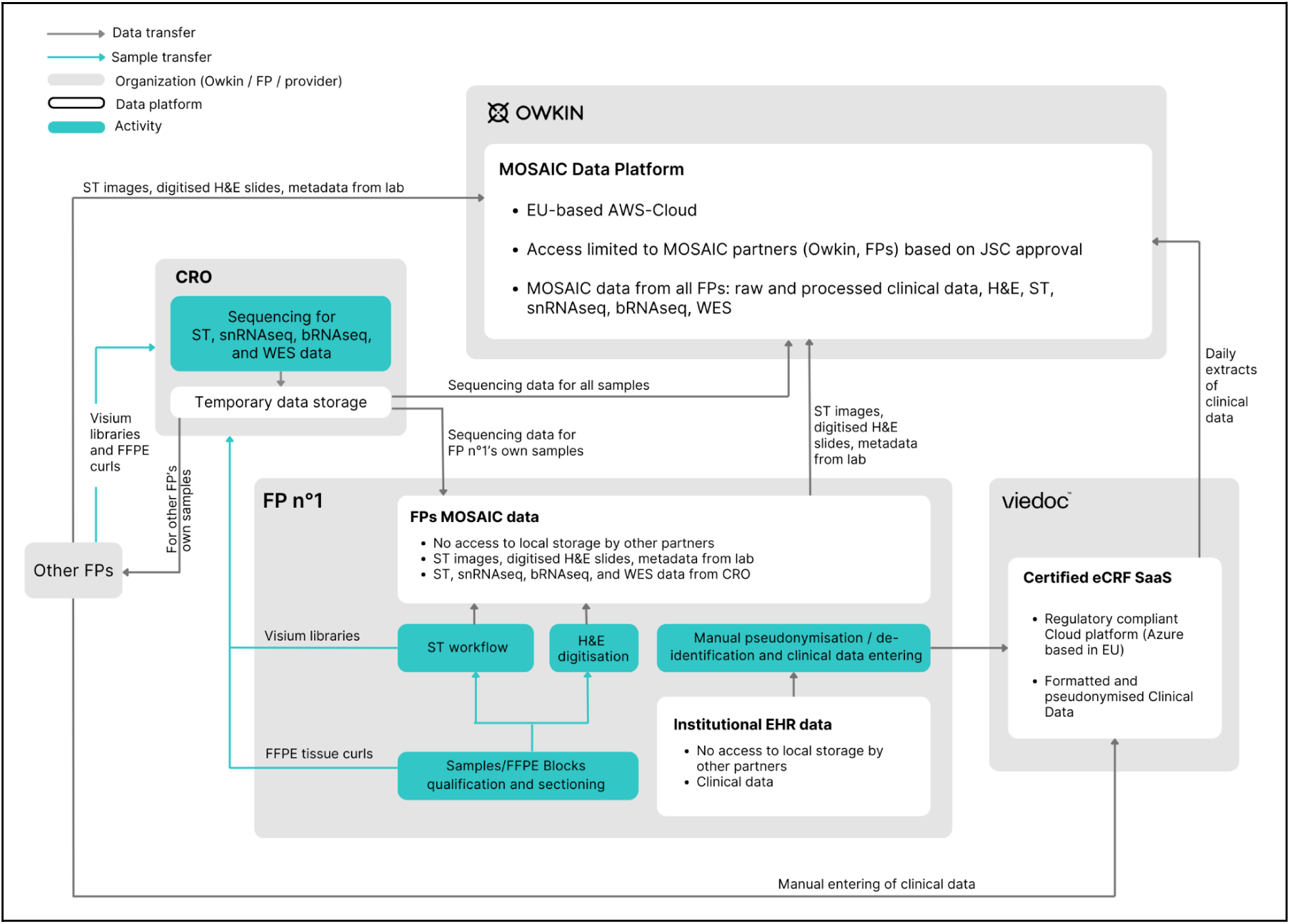
MOSAIC Data Flow and Architecture. The represented diagram is a functional representation of the different blocks involved in data collection, digitization and generation throughout the MOSAIC program lifecycle. The data is generated at the FPs and at the CRO, and it is acquired using a number of component systems integrated into an overall data management system. Data is stored locally at each FP - for its own samples - and centrally on the Owkin platform - for all MOSAIC samples. Viedoc eCRF is the tool used to collect, store and transfer clinical data to Owkin platform. Samples are shipped from FP labs to the CROs where sequencing data is generated and then transferred to the FPs and Owkin. Abbreviations: EHR, electronic health records; FPs, founding partners; eCRF, electronic case report form; Saas, software as a service; H&E, Hematoxylin and Eosin microscopic images; ST, Spatial transcriptomics; snRNAseq, Single Nuclei RNA transcriptomics; bRNAseq, bulk Ribonucleic Acid Sequencing; WES, bulk Whole Exome Sequencing.

**Supplementary Figure 2.**
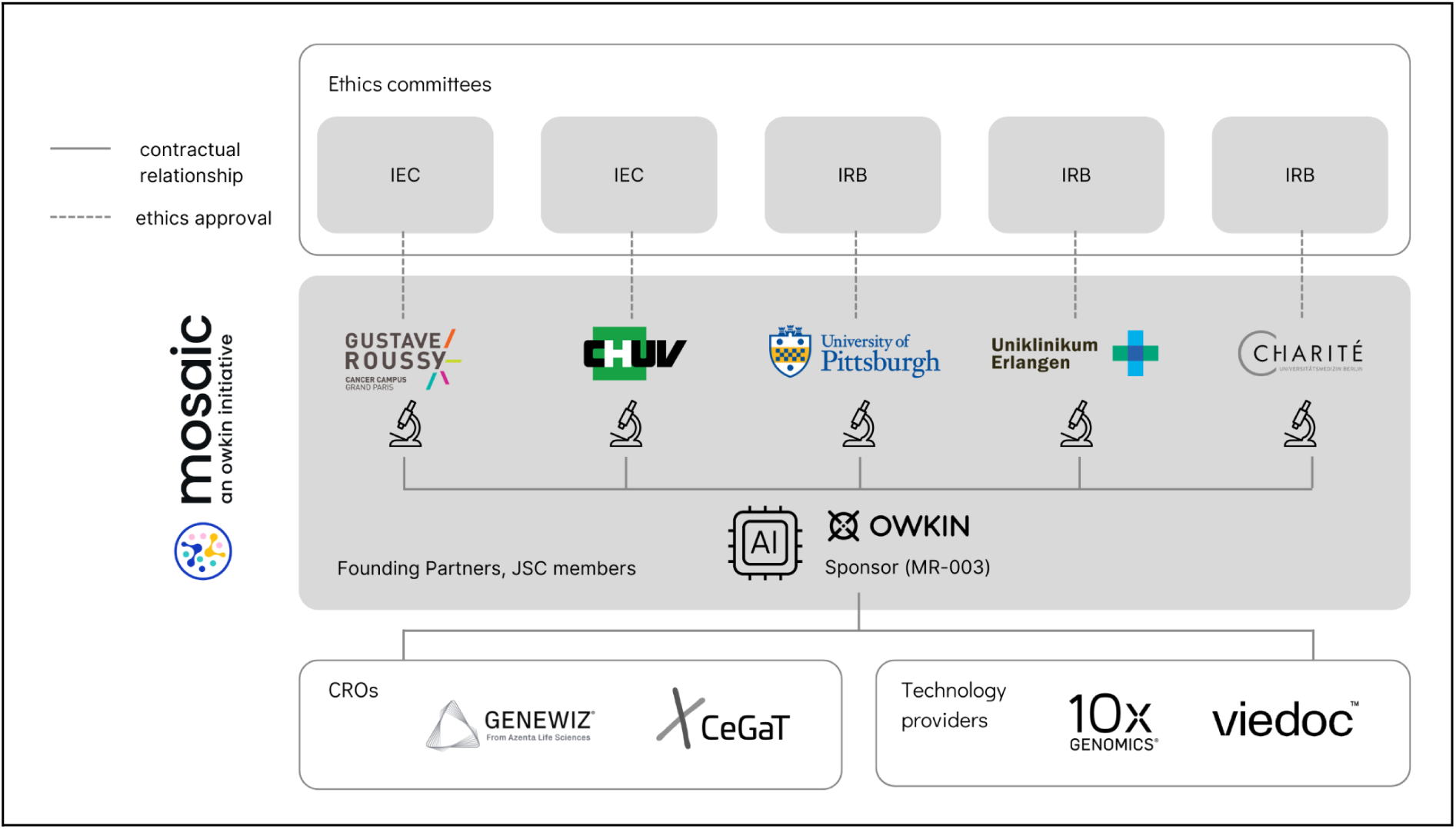
Consortium Governance, Contractual Relationships, and Ethical Oversight Framework. This diagram illustrates the organizational structure and collaborative framework of the MOSAIC consortium. Key contractual relationships between participating institutions and Owkin are depicted, highlighting the formal agreements that facilitate data sharing, resource allocation, and joint project management. Abbreviations: IEC, Institutional Ethics Committees; IRB, Institutional Review Board; JSC, join-steering committee; CROs, contract-research organizations.

**Supplementary Figure 3.**
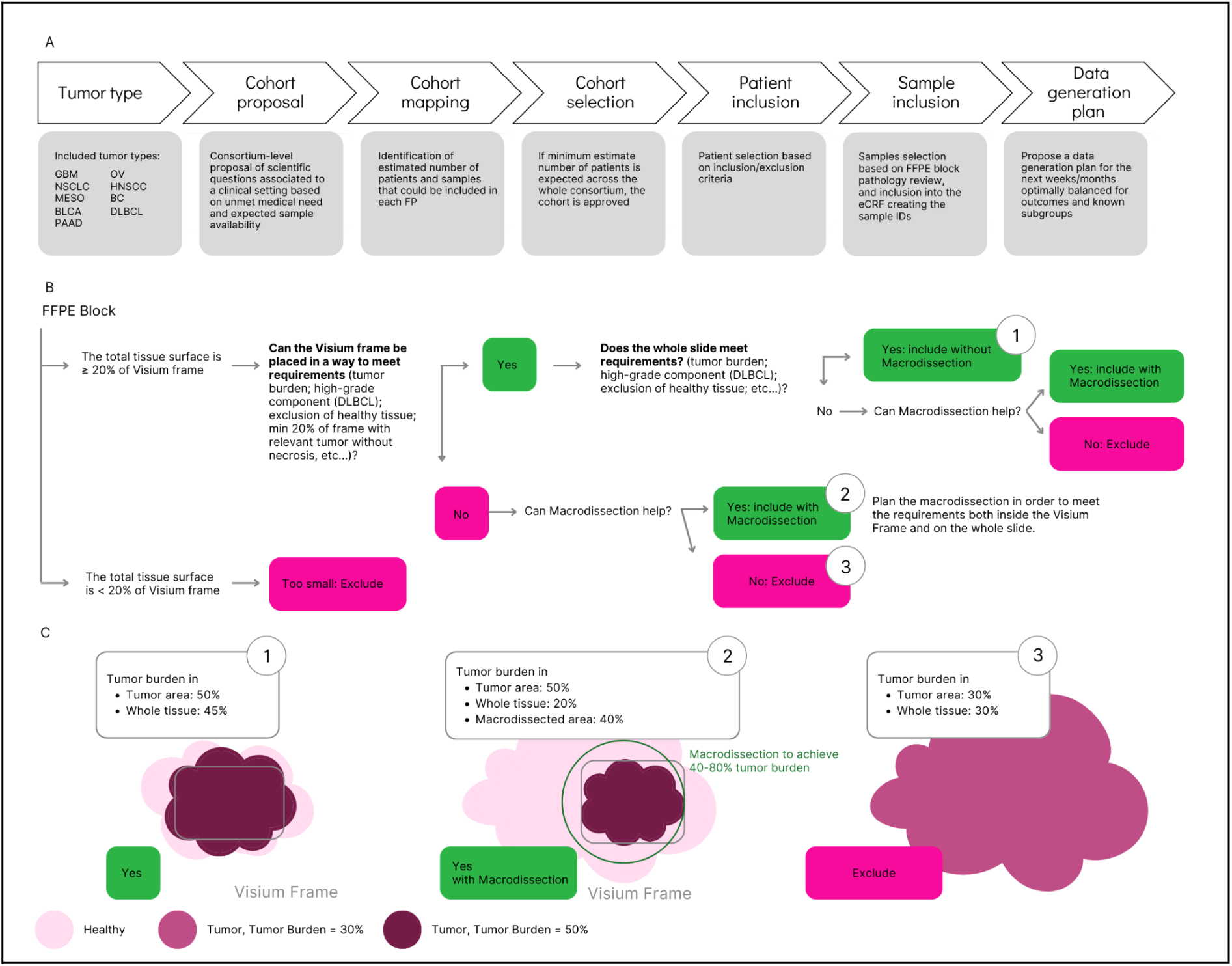
Integrated Workflow from Therapeutic Area Definition to Data Generation and Sample Quality Assurance. (A) Cohort Development and Data Generation Planning: This panel outlines the strategic process for cohort creation and subsequent data generation. The workflow begins with the definition of a tumor type, which informs the formulation of a cohort proposal. Through detailed mapping and selection procedures, candidate patients and samples are identified and evaluated based on established inclusion criteria. The process culminates in the formal inclusion of selected patients and samples into the study, followed by the development of a comprehensive data generation plan. (B) FFPE Block Quality Assurance: FFPE blocks, along with the corresponding archival H&E slides, are sent to the pathology laboratory to ensure that the samples meet the required specifications. Assessment of these criteria was done in a two step manner, first in the 6.5 x 6.5 mm area selected for the Visium frame; second on the whole slide (unless the sample was entirely included in the Visium frame area), relevant for bulkRNAseq, single-nuclei RNAseq, and Whole-Exome Sequencing. (C) The need for macrodissection was adapted accordingly. Abbreviations: FP, founding partner; eCRF, electronic case report form; FFPE, formalin-fixed paraffin-embedded; DLBCL, Diffuse-large B-cell Lymphoma.

**Supplementary Figure 4.**
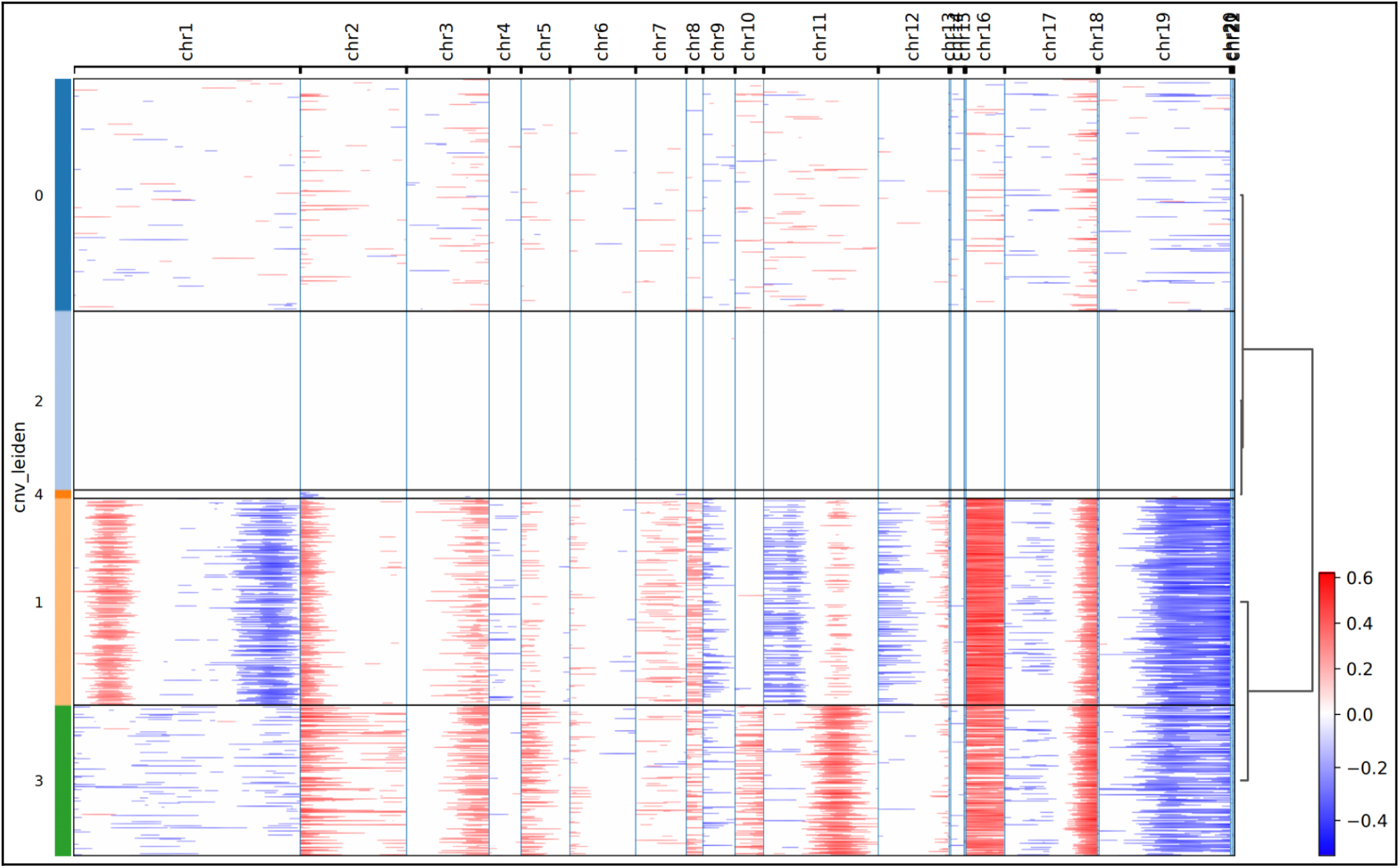
Copy Number Variation inference Shows Different genomic landscape of cancer cell within Case #1 Sample. This figure represents the result of inferCNV form the spot level inference for sample MW_M_051a. In this heatmap, each line is a spot and each column a genomic window of 500 genes. The values indicate the inferred status: red for a gain and blue for a loss of copy-number. All spots (both reference and observation) are included in this representation. Left color annotations indicate the result of the leiden clustering performed on these vector of copy-number inferences per genomic window.

**Supplementary Figure 5.**
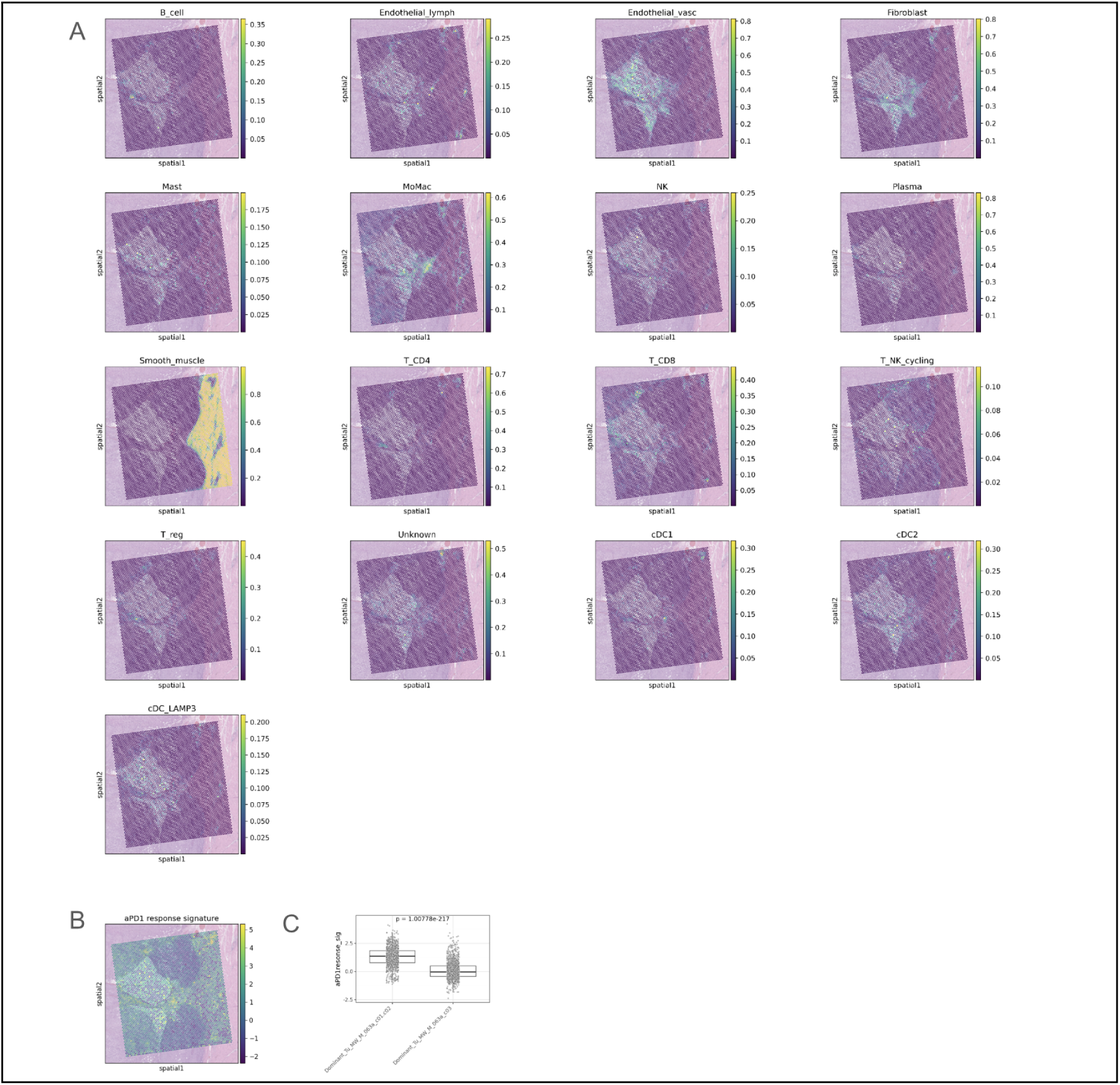
Deconvolution analysis highlighted that the two tumor regions of Case #1 are infiltrated by different non-malignant cell populations and differentially activating aPD1 response signature. A. This figure indicates the cellular fraction heatmaps of non-tumoral cell populations obtained from spot-level deconvolution in sample MW_M_051a. B. Heatmap of the aPD1 response signature activity. C. Boxplot showing the activity of the aPD1 response signature in spots with a dominant fraction of either c01/c02 or c3 cell populations. P-values were calculated with Mann-Whitney statistical test.

**Supplementary Figure 6.**
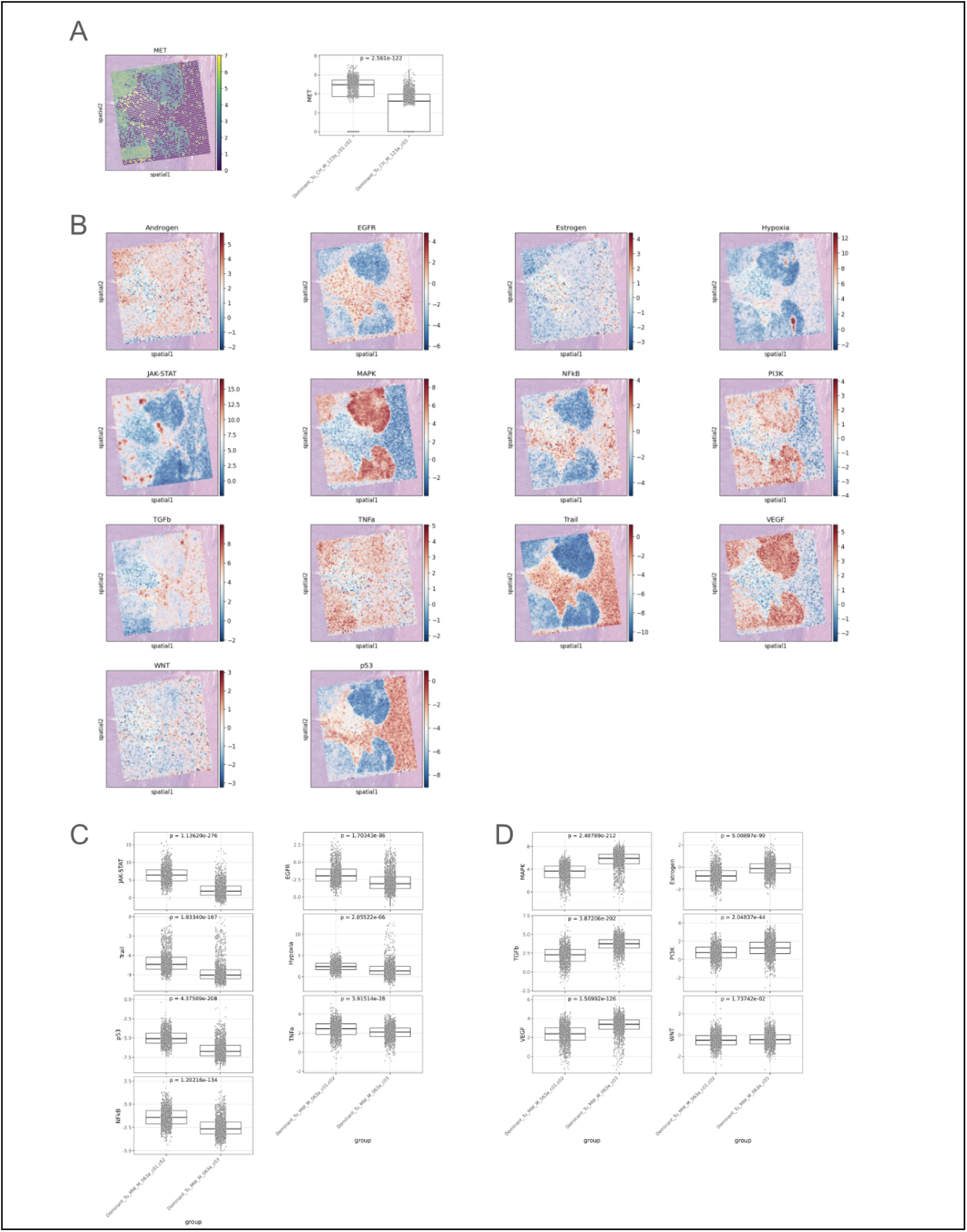
Differential expression and pathway enrichment analysis highlight presence of distinct tumor phenotypes. For patient MW_M_051a, A. The left panel is showing the log1p CPM expression of MET gene while the right panel is indicating, for two sets of patients dominated by c01/c02 or c03 subset fractions, the distribution of met expression. P-value was calculated using Mann-Whitney statistical test. B panel indicates the results of spot level PROGENy pathways activity analysis. C and D shows a boxplot of spots with a dominant fraction of either c01/c02 or c3 cell populations from most significantly different PROGENy pathway activity between the two cancer cell populations. Panel C focuses on pathways activated differentially in c01/c02 while panel D focuses on pathways more active in c03. P-values were calculated with Mann-Whitney statistical test.

**Supplementary Figure 7:**
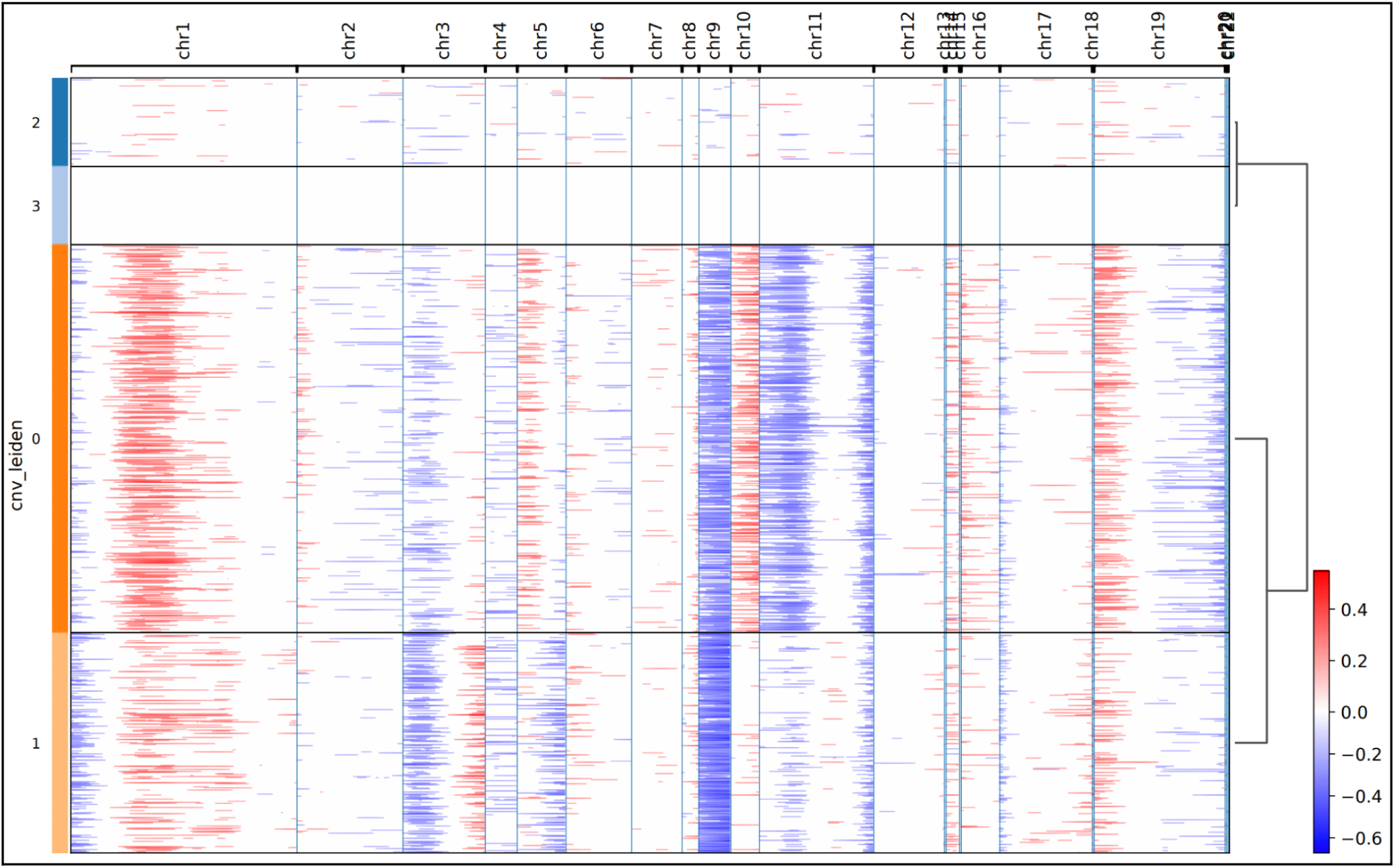
Copy Number Variation inference Shows Different genomic landscape of cancer cell within Case #2 Sample. This figure represents the result of inferCNV from the spot level inference for sample MW_B_079a. In this heatmap, each line is a spot and each column a genomic window of 500 genes. The values indicate the inferred status: red for a gain and blue for a loss of copy-number. All spots (both reference and observation) are included in this representation. Left color annotations indicate the result of the leiden clustering performed on these vectors of copy-number inferences per genomic window.

**Supplementary Figure 8.**
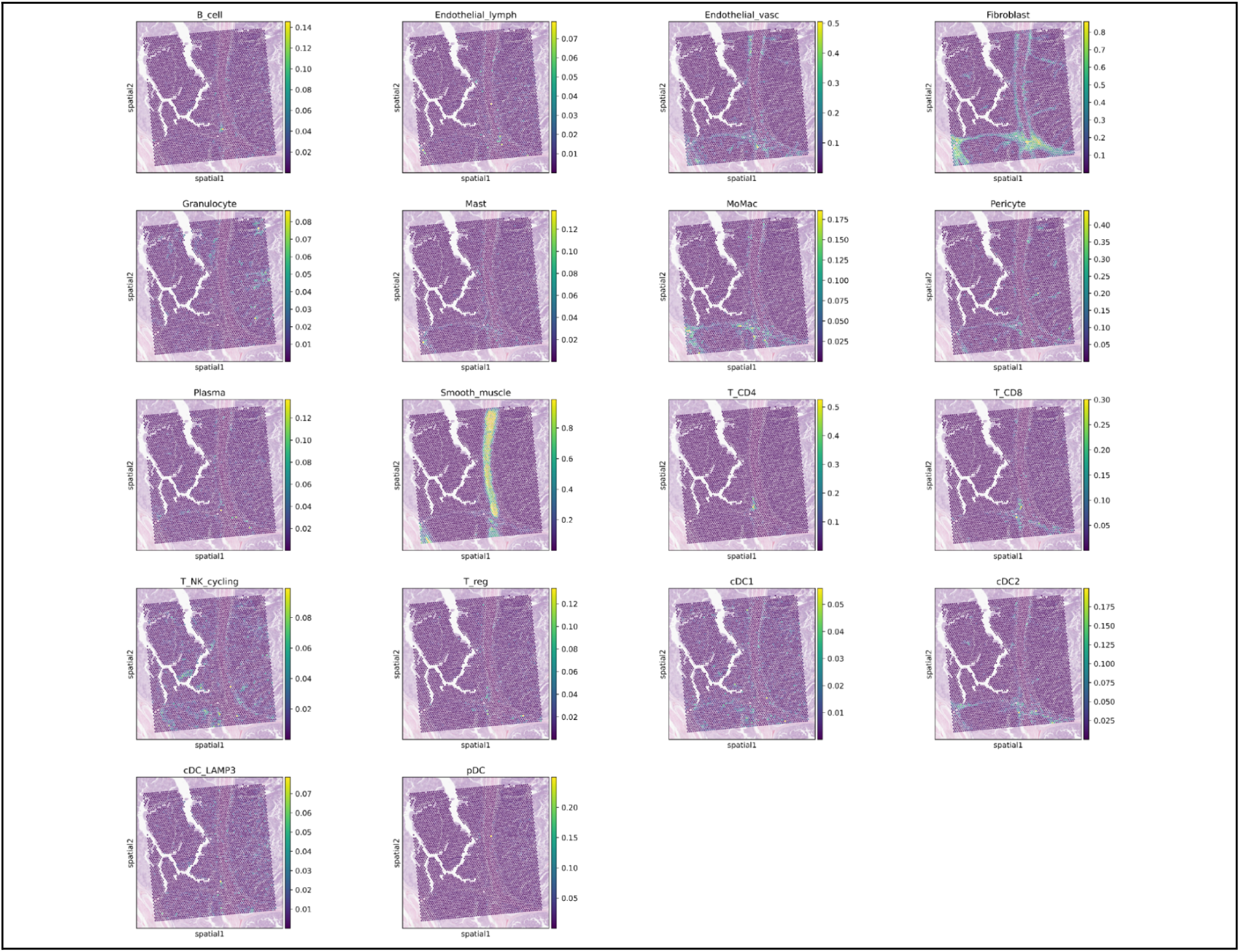
Deconvolution analysis revealed different infiltration of immune cells in Case #2. This figure indicates the cellular fraction heatmaps of non-tumoral cell populations obtained from spot-level deconvolution in sample MW_B_079a

**Supplementary Figure 9.**
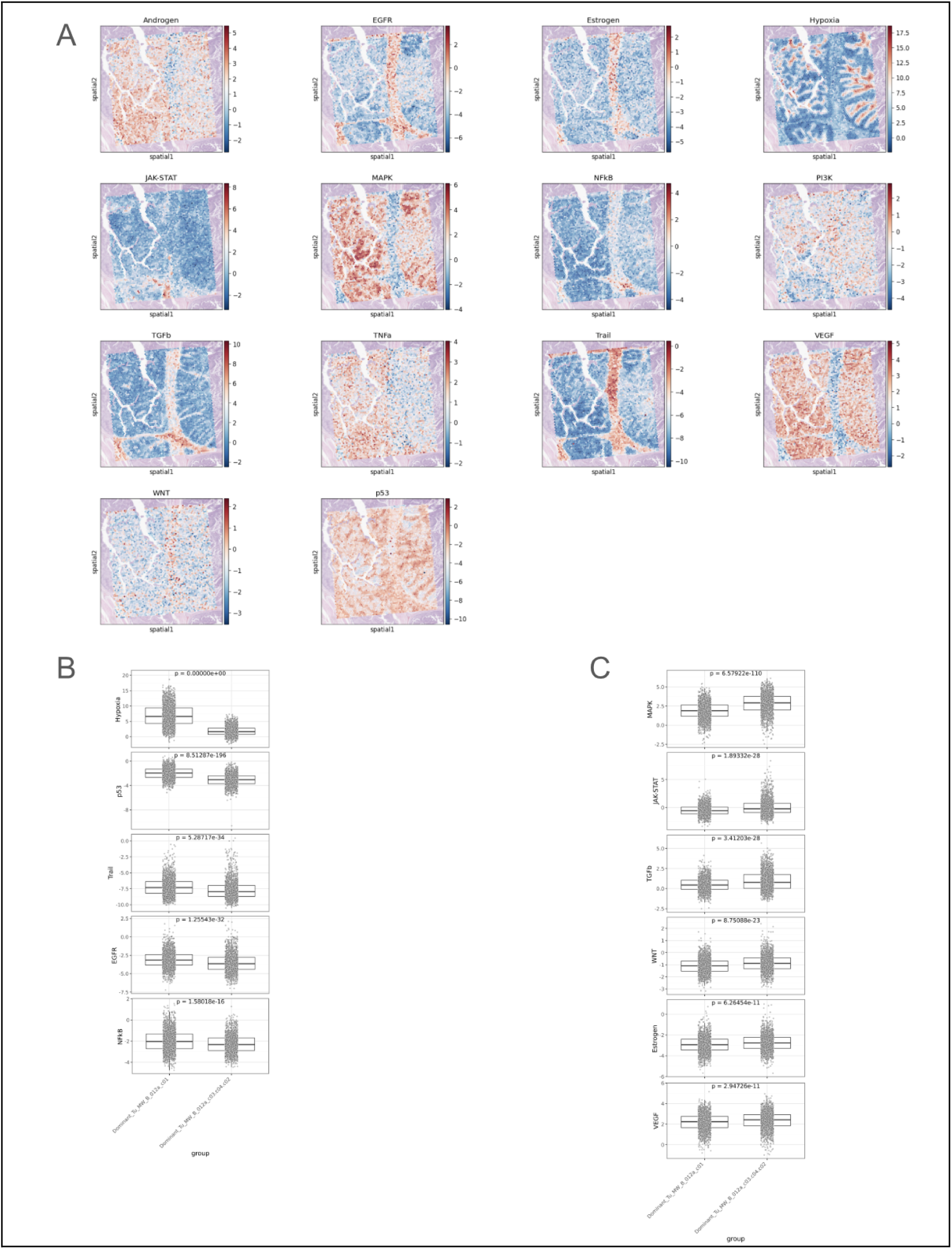
Pathway enrichment analysis highlight presence of distinct tumor phenotypes within Case #3. For patient MW_B_079a, A panel indicates the results of spot level PROGENy pathways activity analysis. B and C show a boxplot of spots with a dominant fraction of either c01/c02 or c3 cell populations from most significantly different PROGENy pathway activity between the two cancer cell populations. Panel C focuses on pathways activated differentially in c01 while panel D focuses on pathways more active in c02/c03/c04. P-values were calculated with Mann-Whitney statistical test.

**Supplementary Figure 10.**
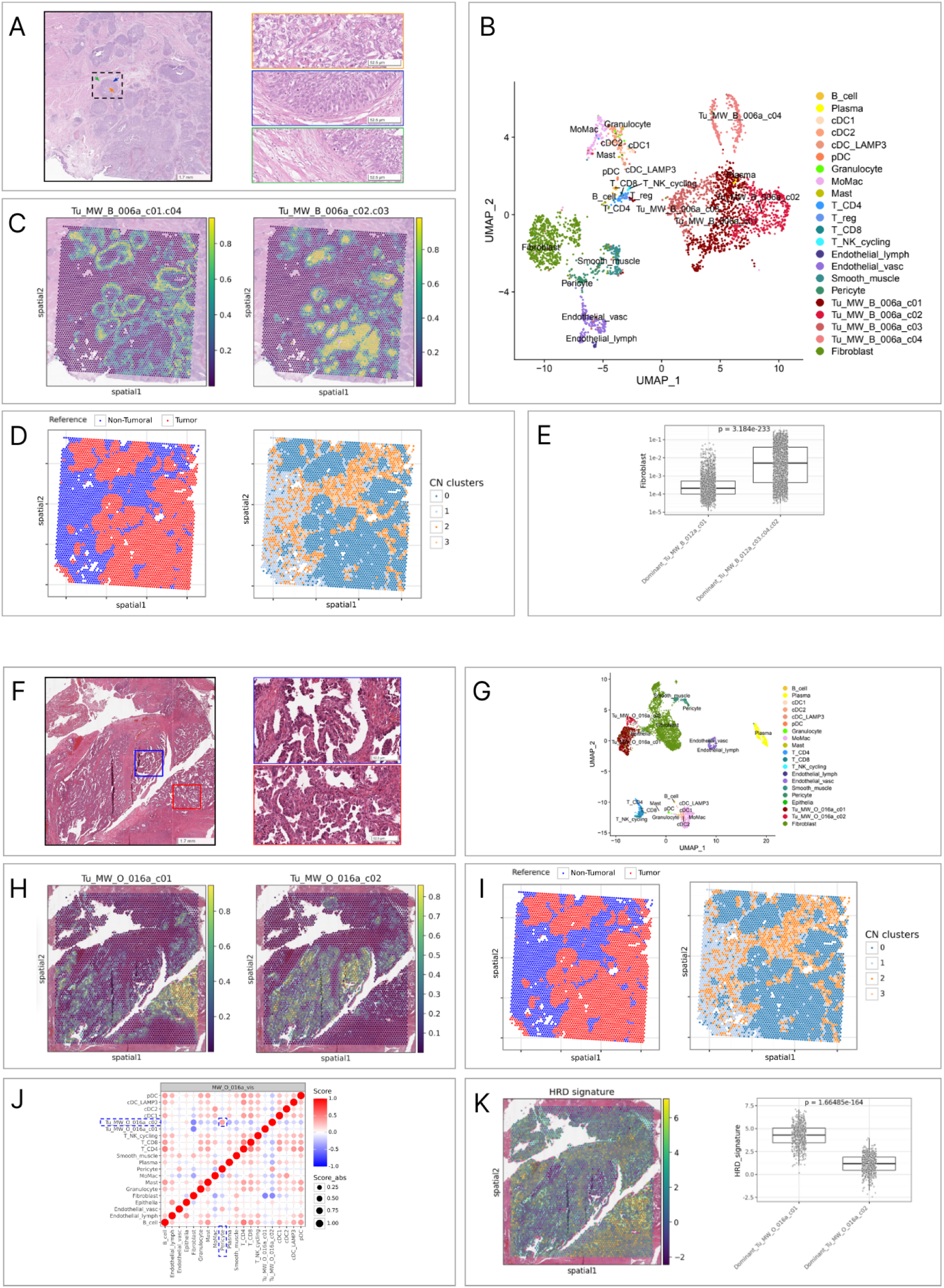
Multimodal integration highlighting intra-tumor heterogeneity in bladder and ovarian tumor cases (Case #3 and Case #4) A. H&E image from a bladder cancer patient (MW_B_006a) showing a tumor “nestˮ architecture. On the right, we can see a detailed view of the tumor nest, which highlights two different cell types. Inside the nest (central area, orange square), round cells with clear cytoplasm and no evident organization are observed. In the peripheral area of the nest, the cells appear elongated and are arranged vertically, in a “peripheral palisading” (outside area, blue square). Finally, surrounding the nest, a thick band of fibroblasts and immune cells encases the structure (green square). B. UMAP embedding from single cell experiment for sample MW_B_006a, cell color is based on the annotation of cell populations. Malignant cell cluster grouping is indicated by dotted line based on their colocalization (see Methods) C. Visium spot level deconvolution of malignant cell populations identified in single-cell for sample MW_B_006a. The two panels are corresponding to spatially distinct tumor cell populations, dotted lines corresponding to two identified areas in panel A. D. The left panel shows the spots used as reference for inferCNV in blue and observation in red, using a 10% tumor fraction threshold. On the right, the spots are colored based on the leiden clustering performed based on inferCNV results, displaying distinct regions based on their CN profiles. E. Boxplot indicating for two sets of patients dominated by c02/c03 or c01/c04 subsets fractions, the distribution of the fibroblast fraction obtained from deconvolution. P-value was calculated using Mann-Whitney statistical test. F. Image of an H&E image from a HGSC tumor (MW_O_016a) showing a papillary pattern in blue and red rectangles. Some minor cytological differences can be observed, but structure is the same at histological level. Panels G, H and I correspond to panels B, C and D for patient MW_O_016. J. Bubble-plot showing the spot-wise Pearson correlation between each pair of cell populations in sample MW_O_016a. Point size corresponds to the absolute value of Pearson correlation while color is indicating the direction of the correlation. Bubbles and cell populations in dashed line squares are the colocalizations specific to one of the two subgroups. K. The left panel is showing the activity of HRD signatures obtained from Guang Peng et al. 2014. The right panel is indicating, for two sets of patients dominated by c01 or c02 subset fractions, the distribution of this signature activity. P-value was calculated using Mann-Whitney statistical test.

**Supplementary Figure 11.**
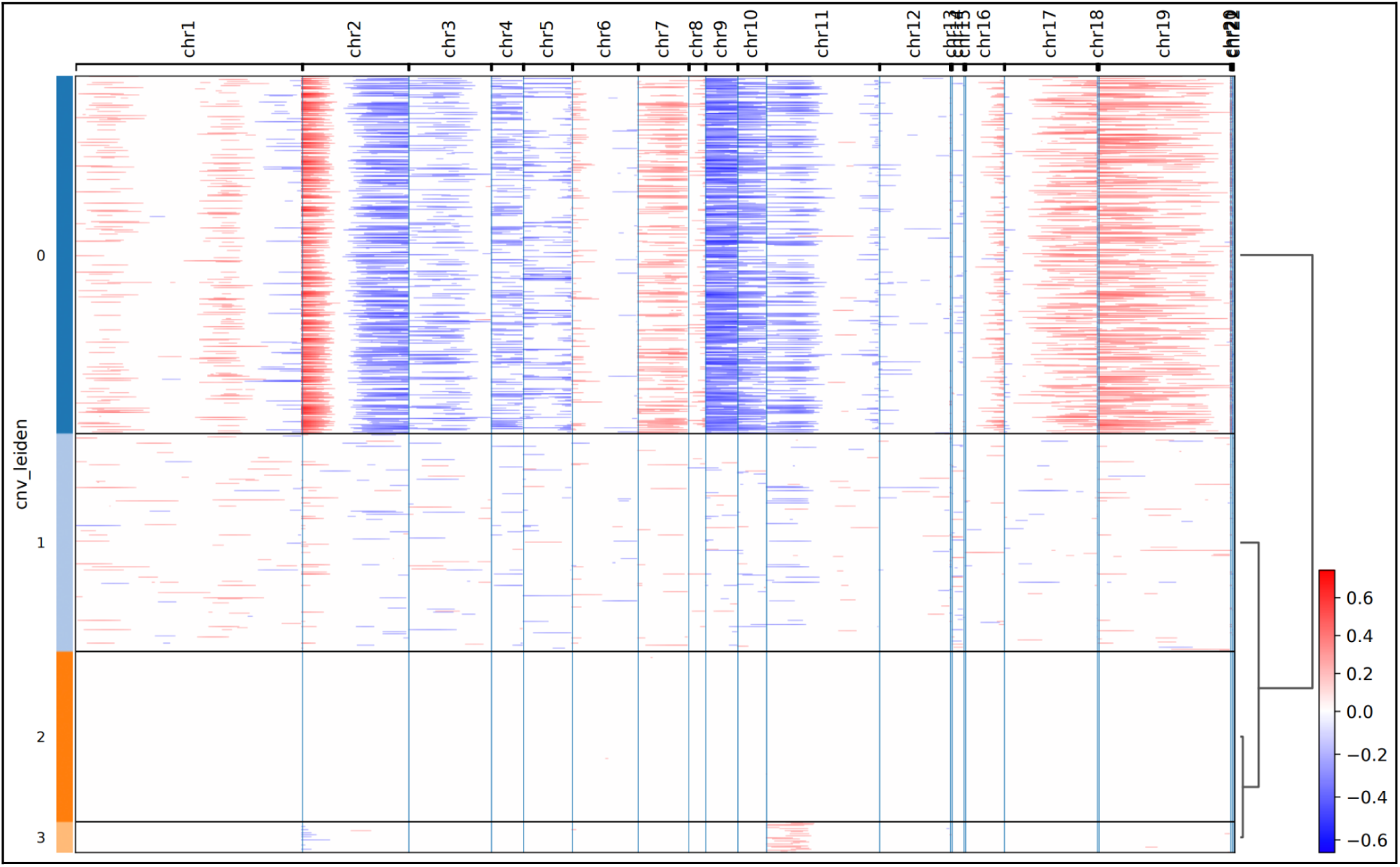
Copy Number Variation inference Shows Different genomic landscape of cancer cell within Case #3. This figure represents the result of inferCNV from the spot level inference for sample MW_B_006a. In this heatmap, each line is a spot and each column a genomic window of 500 genes. The values indicate the inferred status: red for a gain and blue for a loss of copy-number. All spots (both reference and observation) are included in this representation. Left color annotations indicate the result of the leiden clustering performed on these vectors of copy-number inferences per genomic window.

**Supplementary Figure 12.**
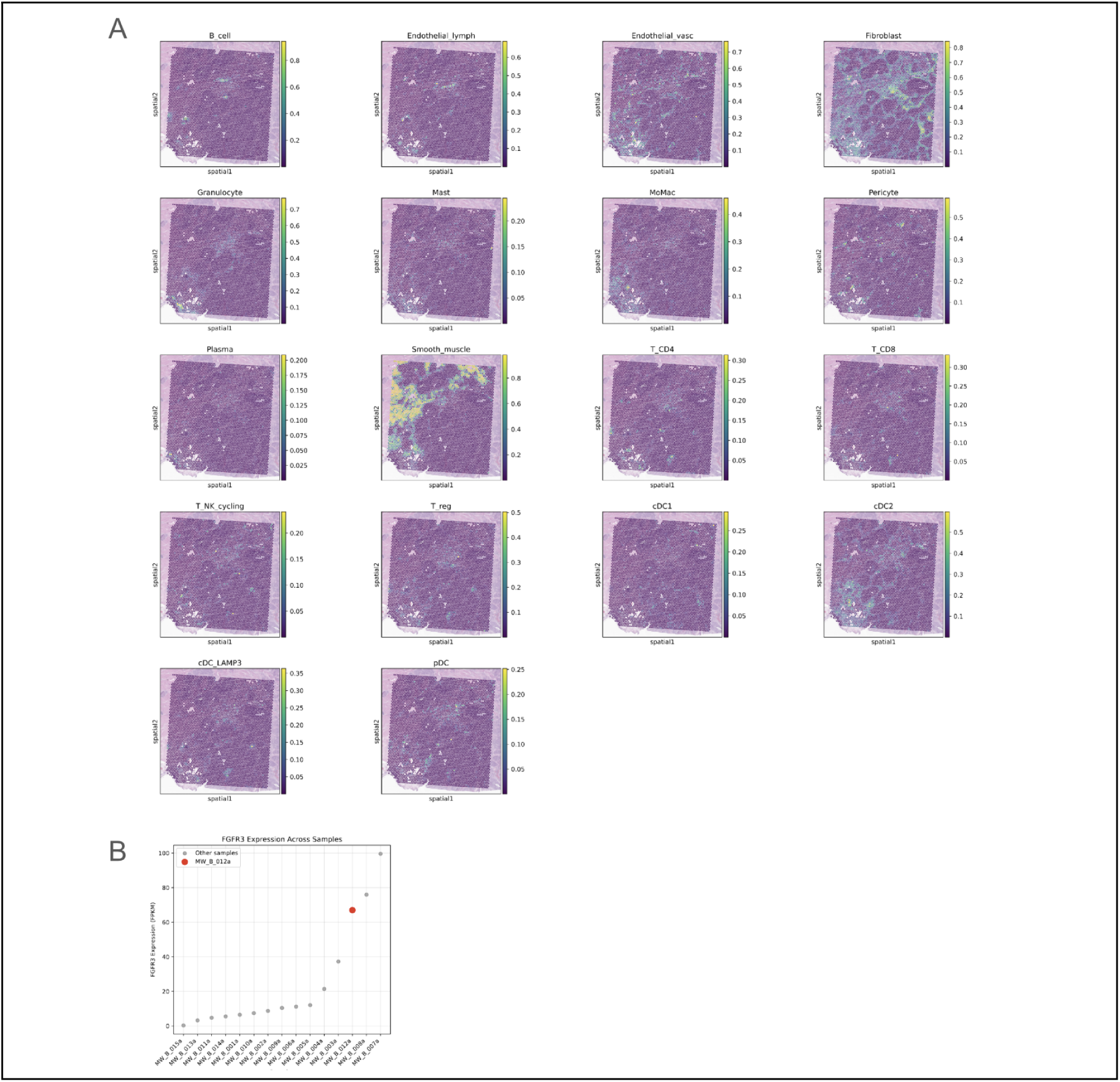
For Case #3 deconvolution analysis confirms insights generated by the histopathological inspection. A. This figure indicates the cellular fraction heatmaps of non-tumoral cell populations obtained from spot-level deconvolution in sample MW_B_006a. B. Scatterplot showing the distribution of FGFR3 FPKM expression in bladder samples of MOSAIC Window cohort, sample MW_B_021a is among the top expressor of FGFR3.

**Supplementary Figure 13.**
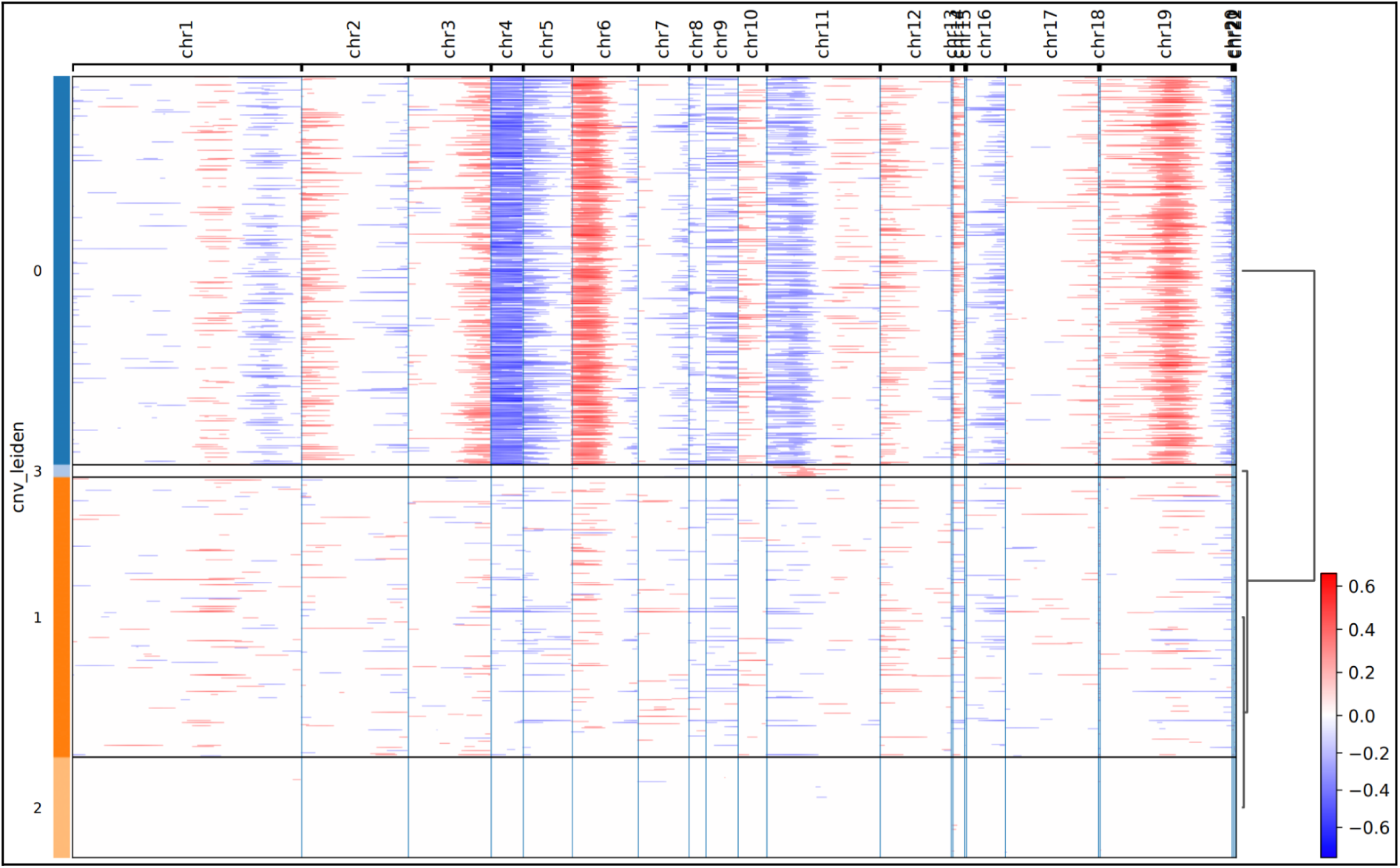
Copy Number Variation inference Shows Different genomic landscape of cancer cell within Case #4. This figure represents the result of inferCNV from the spot level inference for sample MW_O_016a. In this heatmap, each line is a spot and each column a genomic window of 500 genes. The values indicate the inferred status: red for a gain and blue for a loss of copy-number. All spots (both reference and observation) are included in this representation. Left color annotations indicate the result of the leiden clustering performed on these vectors of copy-number inferences per genomic window.

**Supplementary Figure 14.**
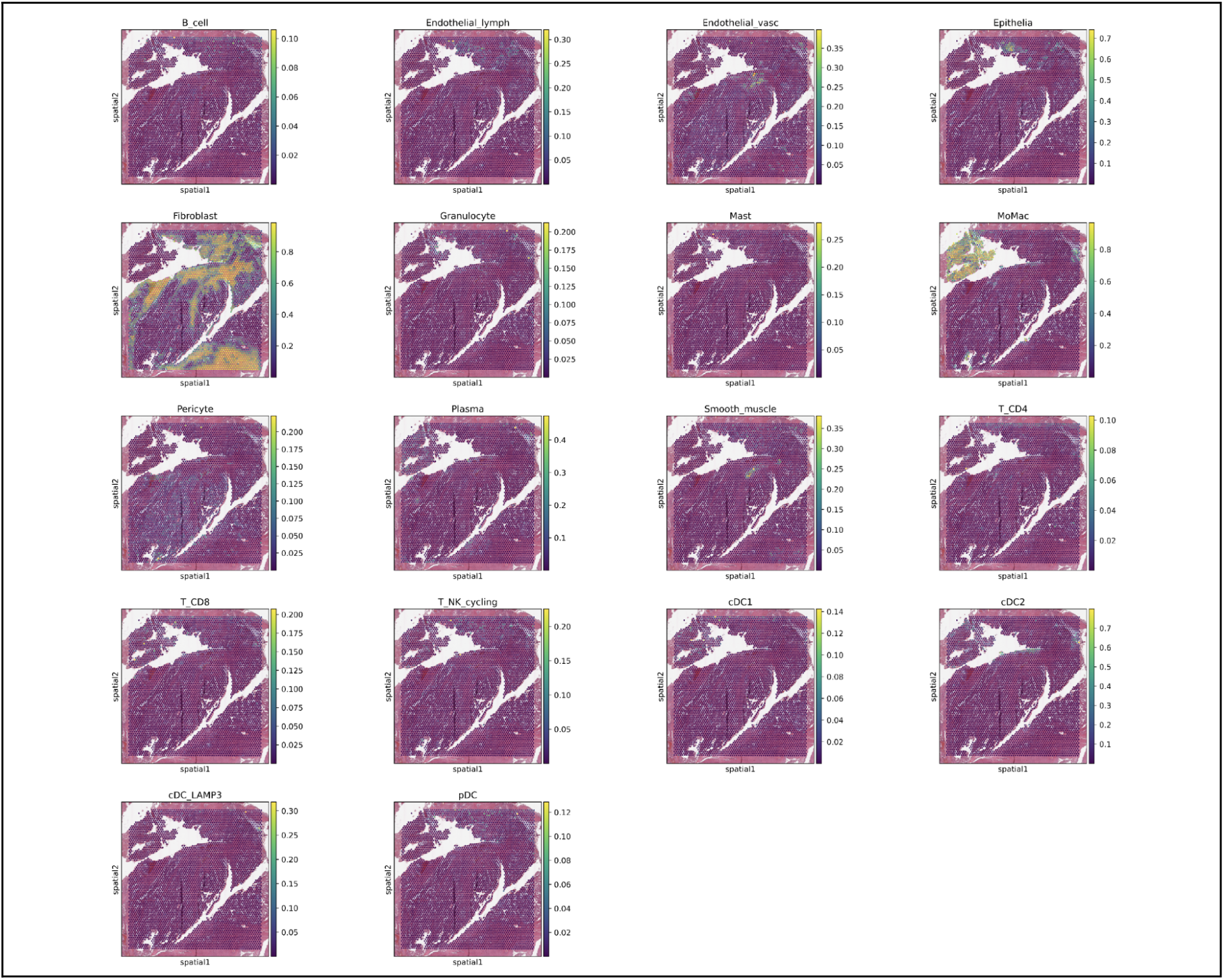
For Case #4 cell deconvolution analysis suggests that the c02 subset is strongly associated with pericytes. This figure indicates the cellular fraction heatmaps of non-tumoral cell populations obtained from spot-level deconvolution in sample MW_O_016a

**Supplementary Figure 15.**
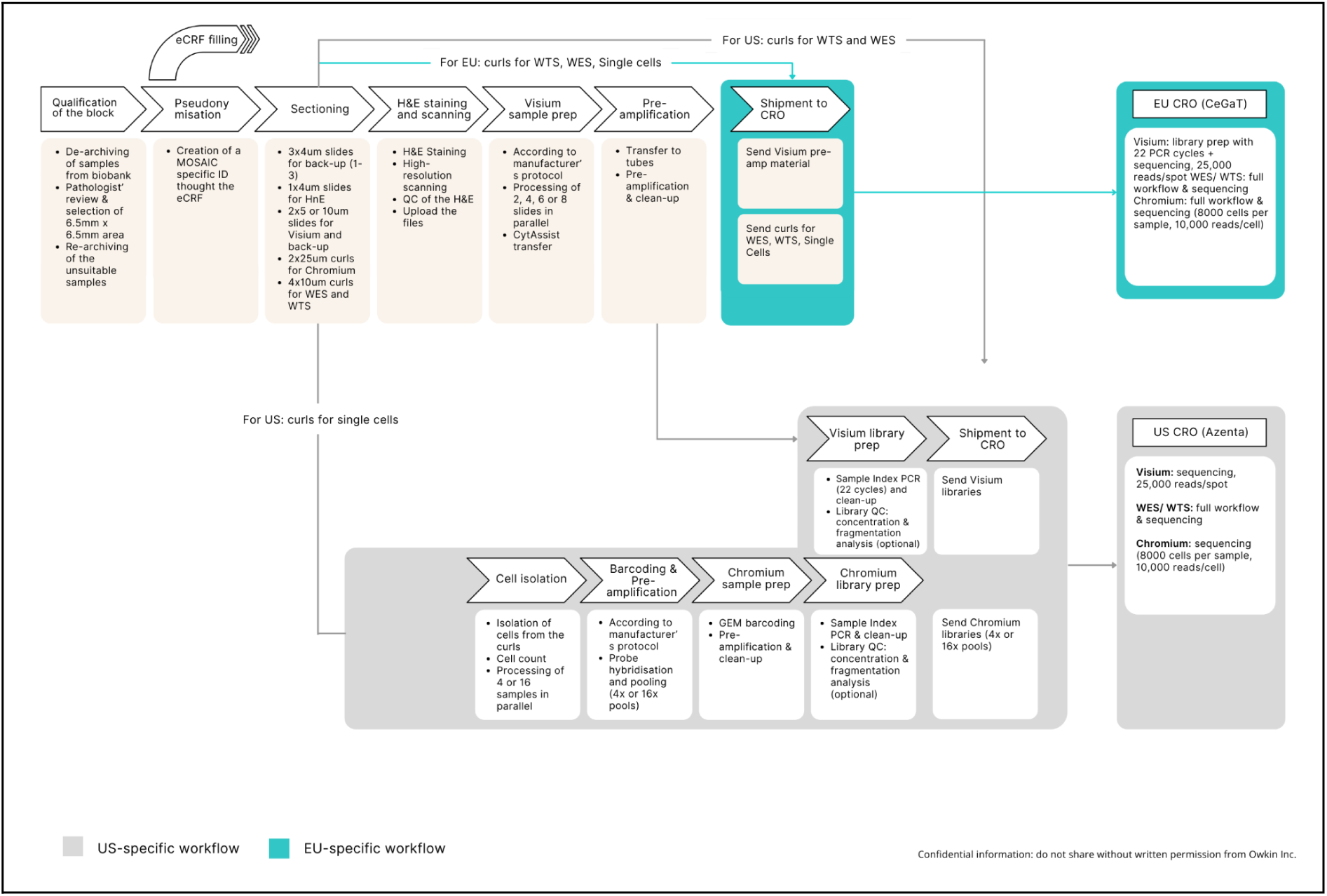
Comprehensive Overview of the Laboratory Workflow. This schematic illustrates the stepwise laboratory workflow for processing tissue samples prior to multi-omics analyses. The Visium sections are processed up to either the pre-amplification step in the European centers or the final library preparation stage at the University of Pittsburgh. Chromium workflow in the US is performed until the library preparation. Subsequently, the libraries are shipped to the CRO along with the curls designated for bulk RNAseq and Whole-Exome Sequencing for further processing and sequencing. Abbreviations: eCRF, electronic case report form; WTS, whole-transcriptome sequencing; WES, whole-exome sequencing; CRO, contract research organization; GEM, Gel Beads-in-Emulsion; PCR, polymerase chain reaction; QC, quality control.

